# Expressional Diversity and Cancer-prone Phenotypes in Cigarette-smoking Lungs at Single Cell Resolution

**DOI:** 10.1101/2021.12.09.472029

**Authors:** Jun Nakayama, Yusuke Yamamoto

## Abstract

Single-cell RNA-seq (scRNA-seq) technologies have been broadly utilized to reveal molecular mechanisms of respiratory pathology and physiology at single-cell resolution. Here, we established single-cell meta-analysis (scMeta-analysis) by integrating data from 8 public datasets, including 104 lung scRNA-seq samples with clinicopathological information and designated a cigarette smoking lung atlas. The atlas revealed early carcinogenesis events and defined the alterations of single-cell transcriptomics, cell population, and fundamental properties of biological pathways induced by smoking. In addition, we developed two novel scMeta-analysis methods: VARIED (Visualized Algorithms of Relationships In Expressional Diversity) and AGED (Aging-related Gene Expressional Differences). VARIED analysis revealed expressional diversity associated with smoking carcinogenesis. AGED analysis revealed differences in gene expression related to both aging and smoking states. The scMeta-analysis pave the way to utilize publicly -available scRNA-seq data and provide new insights into the effects of smoking and into cellular diversity in human lungs, at single-cell resolution.

## Introduction

Smoking is the leading risk factor for early death, and its negative effects present individual and public health hazards [1, 2]. Cigarette smoke is a mixture of thousands of chemical compounds generated from tobacco burning [3] that causes chronic airway inflammation, reactive oxygen species (ROS) production, and DNA damage. Specifically, it has been discovered that smoking injures the respiratory organs and cardiovascular system and causes carcinogenesis, chronic obstructive pulmonary disease (COPD), and atherosclerosis [4]. In particular, the incidence of lung squamous carcinoma is significantly increased by cigarette smoking [5, 6].

Single-cell RNA-seq (scRNA-seq) technologies have been broadly utilized to reveal the molecular mechanisms of respiratory diseases and physiology at single-cell resolution. scRNA-seq in human lungs identified novel cell populations and cellular diversity [7-13]. However, there are several concerns regarding scRNA-seq analysis. One of these concerns is sample size, that is, that clinical scRNA-seq analyses could be biased due to insufficient sample sizes. A possible solution is meta-analysis of scRNA-seq data. The recently developed single-cell meta-analysis (scMeta-analysis) method has been considered a powerful tool for large-scale analysis of integrated single-cell cohorts. The scMeta-analysis shows robust statistical significance and the capacity to compare the results among different studies at the single-cell level. In fact, integrated scMeta-analysis of a number of cohorts has revealed a previously unappreciated diversity of cell types and gene expression; for example, scMeta-analysis of lung endothelial cells, including human and mouse datasets, revealed novel endothelial cell populations [14-17]. In addition, comparative analysis of scRNA-seq cohorts revealed pan-cancer tumor-specific myeloid lineages [18].

In this study, we integrated 8 publicly available datasets comprising 104 lung scRNA-seq samples and analyzed a total of 230,890 single cells to construct a cigarette smoking lung atlas. The scMeta-analysis of the cigarette smoking lung atlas defined single-cell gene expression according to smoking, age, and gender. In addition, we developed novel scMeta-analysis methods: VARIED (Visualized Algorithms of Relationships In Expressional Diversity) analysis and AGED (Aging-related Gene Expressional Differences) analysis with clinical metadata. VARIED analysis revealed the diversity of gene expression associated with cancer-related events in each cell population, and AGED analysis revealed the expressional differences in relation to both aging and smoking states.

## Materials and Methods

### scRNA-seq data collection from public databases

The scRNA-seq cohorts were downloaded from the public Gene Expression Omnibus (GEO) and European Genome-Phenome Archive (EGA) databases (Supplementary Table S1). We collected scRNA-seq samples of human lungs for which smoking states information was available. From physiological studies of the lung airway [19], all 10 never-smoker samples were extracted from the EGA00001004082 dataset [20], and 1 never-smoker and 3 smoker samples were extracted from the GSE130148 dataset [13]. From idiopathic pulmonary fibrosis (IPF) studies, 5 never-smoker and 3 smoker samples were extracted from a total of 17 samples in the GSE122960 dataset [21], 1 never-smoker and 7 smoker samples were extracted from a total of 34 samples in the GSE135893 dataset [12], and 22 never-smoker and 23 smoker samples were extracted from a total of 78 samples in the GSE136831 dataset [11]. From studies of lung disease in smokers, 3 never-smoker and 3 smoker samples were extracted from the GSE123405 dataset [22], and 3 never-smoker and 9 smoker samples were extracted from the GSE173896 dataset [23]. From lung cancer studies, 4 never-smoker and 7 smoker samples were extracted from a total of 58 samples in the GSE131907 dataset [24]. A total of 104 samples (never-smoker: 49, smoker: 56) were collected, and the details of the extracted samples are shown in Supplementary Table S2. These datasets were imported into R software version 4.2.0. and transformed into Seurat objects with the package Seurat version 4.3.0 [25]. The Seurat objects from the different datasets were then integrated in R.

### Integration of datasets, data quality control and removal of batch effects

The integrated dataset was subjected to normalization, scaling, and principal component analysis (PCA) with Seurat functions. Removal of low-quality cells was performed against the merged dataset before batch effect removal according to the following criteria (nFeature_RNA > 1000 and percent.mt < 20). The expression counts of each sample were normalized by SCTranscform method version 0.3.5. [26]. Doublet cells in the integrated dataset were removed by DoubletFinder method version 2.0.3. [19, 27]. To remove the batch effect between cohort studies, Harmony version 0.1.1. algorithms were applied to the integrated datasets [28, 29] following the instructions in the Quick start vignettes (https://portals.broadinstitute.org/harmony/articles/quickstart.html).

### Cell type annotation and cell cycle scoring

Clustering of neighboring cells was performed by the functions ‘FindNeighbors’ and ‘FindClusters’ from Seurat using Harmony reduction. First, the clusters were grouped based on the expression of tissue compartment markers (for example, *EPCAM* for epithelia, *CLDN5* for endothelia, *COL1A2* for fibroblasts, and *PTPRC* for immune cells) (Figure 1C and Supplementary Figure S3) and then annotated in detail according to “A molecular cell atlas of the human lung” [7]. Cell cycle analysis was performed with the ‘CellCycleScoring’ function of Seurat.

**Figure 1.**
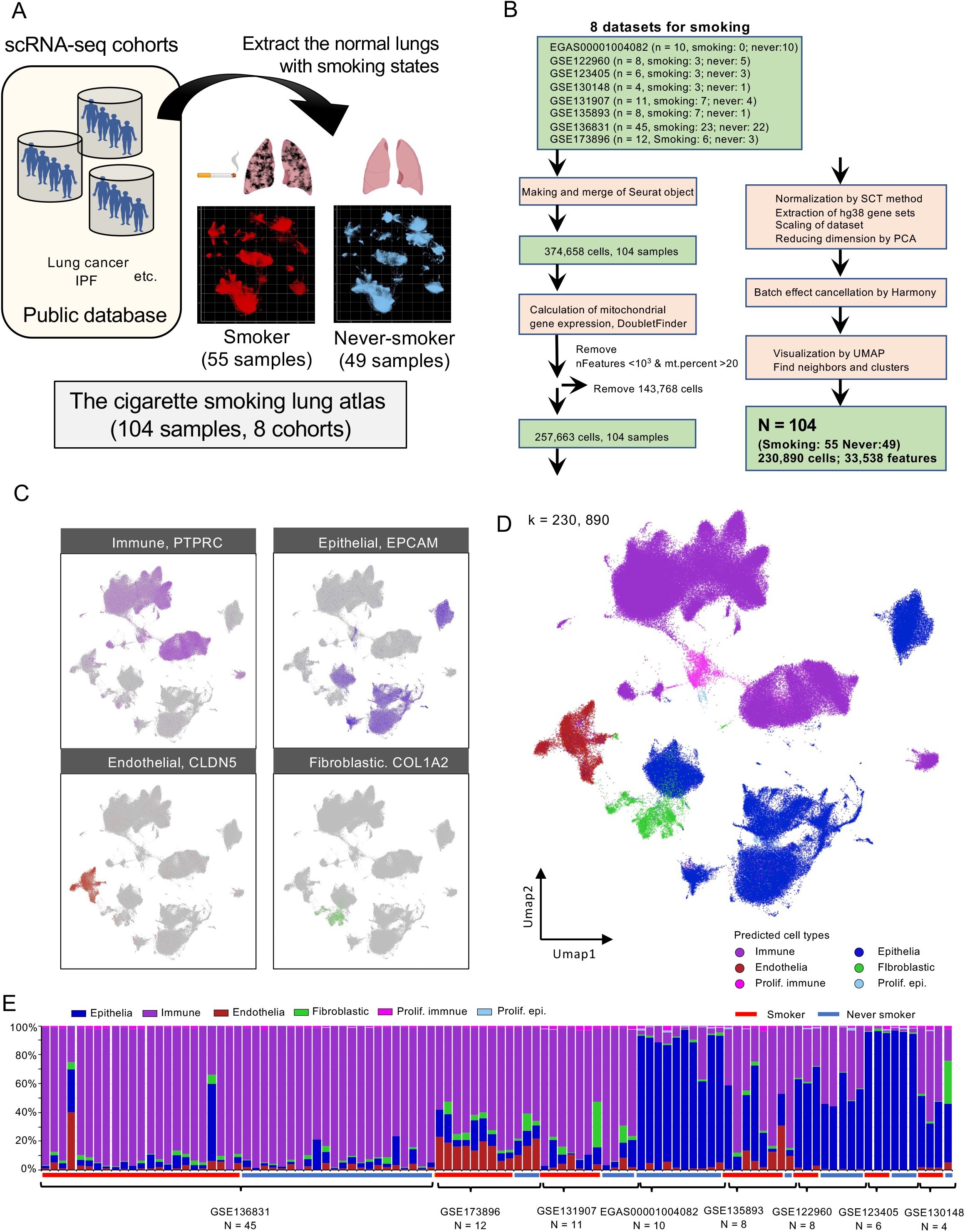
Establishment of the integrated lung dataset with cigarette smoking states from 8 scRNA-seq cohorts. **A.** Overview of the establishment of the integrated lung dataset with cigarette smoking states. The control lung scRNA-seq data from 8 publicly available datasets were obtained and integrated with smoking states information. **B.** Flow diagram of the establishment of the integrated scRNA-seq lung dataset with cigarette smoking. Eight publicly available scRNA-seq datasets were downloaded and combined in Seurat. Doublet cells were removed by DoubletFinder. The datasets were normalized by SCTramsform, integrated by Harmony to adjust for batch effects. **C.** Representative marker expression patterns for the cell type clusters shown in the UMAP plot. **D.** A UMAP plot displaying 230,890 single human lung cells of 55 smokers and 49 never-smokers. Each dot represents a single cell, and cell clusters are classified as immune cells, epithelial cells, endothelial cells, and fibroblasts. **E.** Cell populations of immune cell, epithelial cell, endothelial cell, and fibroblast clusters across the 104 samples. Smokers, 55 cases; never-smokers, 49 cases.

### VARIED (Visualized Algorithms of Relationships In Expressional Diversity) analysis

To evaluate the expressional heterogeneity in the cell populations, we calculated the correlation coefficients for each cell population between smokers and never-smokers. In each cluster, normalized closeness centrality was calculated in R using ggraph version 2.1.0. and igraph version 1.3.5. packages, as previously described [23, 30].

where r is the absolute value of Pearson’s correlational coefficient and n is the number of cells in the cluster.

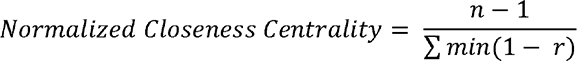

### GSVA scoring and survival analysis

GSVA analysis was performed using RNA-seq dataset of TCGA LUSC cohorts by ‘GSVA’ package version 1.44.5. in R [31]. RNA-seq dataset and clinical information of the lung squamous carcinoma patients were downloaded from TCGA Data Portal [32]. The signature genes of basal-d smoker clusters showed in Supplementary Table S5. Scoring method by ‘gsva’ algorithms was utilized for calculation of enrichment score in the lung squamous cancer patients. We subjected clinical status and gene expression data to survival analysis using ‘survminer’ version 0.4.9. and ‘survival’ packages version 3.4-0 in R. The dataset is available at https://portal.gdc.cancer.gov/.

### Module analysis

Module analysis was performed by the ‘AddModuleScore’ function in Seurat using the gene lists from MSigDB (https://www.gsea-msigdb.org/gsea/msigdb/). The EMT module (HALLMARK_EPITHELIAL_MESENCHYMAL_TRANSITION), heme metabolism module (HALLMARK_HEME_METABOLISM), ROS module (HOUSTIS_ROS), autophagy module (REACTOME_AUTOPHAGY), IFN signaling module (REACTOME_INTERFERON_SIGNALING), senescence module (REACTOME_CELLULAR_SENESCENCE), circadian module (REACTOME_CIRCADIAN_CLOCK), mitophagy module (REACTOME_MITOPHAGY), pyroptosis module (REACTOME_PYROPTOSIS), and ferroptosis module (WP_FERROPTOSIS) were subjected to module analysis in each cell population.

### Pathway enrichment analyses and IPA

We performed enrichment analysis against the marker gene list in each cluster between male and female smokers by the ‘ClusterProfiler’ version 4.4.4. [33] and ‘ReactomePA’ version 1.40.10. [34] packages in R. Gene symbols were converted to ENTREZ IDs using the ‘org.Hs.eg.db’ package version 3.10.0. Pathway datasets were downloaded from the Reactome database. Pathway enrichment analysis using the ‘enrichPathway’ function was performed by the BH method. Marker genes of the basal-px cluster in smokers and never-smokers were calculated by ‘FindMarkers’ with the MAST method [35]. Enrichment analysis of basal-px was performed using QIAGEN Ingenuity Pathway Analysis software.

### AGED (Aging-related Gene Expressional Differences) analysis

We calculated the average expression of all genes in each cluster in both smokers and never-smokers and performed regression analysis in correlation with gene expression and patient age by R. Next, we calculated the differences in slopes (delta) in smokers and never-smokers via regression analysis and extracted the genes with the highest delta to be shown in a heatmap.

### Code and data availability

The datasets GSE122960, GSE123405, GSE130148, GSE131907, GSE135893, GSE136831, and GSE173896 are available in the NCBI GEO database (https://www.ncbi.nlm.nih.gov/geo/). The EGAS00001004082 dataset is available in the EGA database (https://ega-archive.org/). The source code of scMeta-analysis and integrated datasets is available on GitHub (https://github.com/JunNakayama/scMeta-analysis-of-cigarette-smoking).

### Data visualization

The dimensionality-reduced cell clustering is shown as a UMAP plot by the function ‘runUMAP’. Heatmaps were drawn by Morpheus from the Broad Institute. A ridge plot was drawn using the ‘ggridges’ version 0.5.4. package in R. Violin plots were drawn using the ‘ggplot2’ version 3.4.0. package in R.

### Statistical Analysis

Correlation coefficients were calculated by Spearman correlation in R. Welch’s t test or Tukey’s or Dunnett’s multiple comparison test was used for comparison of the datasets. Log-rank test was used for survival analysis in R. Significance was defined as P < 0.05.

## Results

### Establishment of Integrated single-cell lung dataset with cigarette smoking

According to scRNA-seq collection criteria (see methods), we chose 8 publicly available datasets of lung scRNA-seq data to construct a cigarette smoking lung atlas (Figure 1A). To this end, we collected data from 374,658 single cells from 104 scRNA-seq samples (smoker: 55 samples, never-smoker: 49 samples, Figure 1A). In the process of quality control with Seurat in R, 143,768 low-quality single cells (nFeatures < 10^3^ & mt.percent > 20%) were removed. Doublet cells were removed identified by DoubletFinder algorithm [27]. The single cells were normalized by SCTransform method in Seurat [26]. Integration of the 8 datasets was performed by the Harmony algorithm with the smoking states of scRNA-seq samples [28] (Figure 1B). Integrated single-cell transcriptome data were linked with clinical metadata such as smoking states, age, gender, and race (Supplementary Table S1, S2, and Supplementary Figure S1A). The normalized dataset was integrated for removal of batch effect by Harmony algorisms (Supplementary Figure S1B). The cigarette smoking lung atlas is composed of a total of 230,890 single cells. The density plot showed that the majority of single cells in the atlas were immune cells and epithelial cells (Supplementary Figure S1C). UMAP plots with cell type-specific markers (*PTPRC* as an immune marker, *EPCAM* as an epithelial marker, *CLDN5* as an endothelial marker, and *COL1A2* as a fibroblast marker) showed an obvious segregation of immune, epithelial, endothelial, and fibroblastic lineages (Figure 1C). There were 118,364 single cells in the smoker group and 112,526 single cells in the never-smoker group (Supplementary Figure S2A). Comparison of the atlases by smoking states revealed that most of the cell populations in the UMAP plot overlapped; however, parts of epithelial clusters were specific to the never-smoker group (Supplementary Figure S2A). To confirm that the integration of the 8 datasets reduced bias, we showed the atlas marked with the datasets (Figure 1D). All major clusters seemed to overlap among the 8 datasets (Supplementary Figure S2B), although the populations of cells were different in each dataset (Figure 1E). This difference in cell populations could be caused by differences in tissue collection and cell isolation processes.

In the atlas with all cell types (Figure 1D), we first identified the cell types present within the atlas according to the lung cell markers in the human lung scRNA-seq atlas [7] (Supplementary Figure S3). To investigate the cell types in further detail, we extracted subsets of “epithelia” (Figure 2A), “fibroblasts” (Figure 2B), “endothelia” (Figure 2C), “lymphoids” (Figure 2D), and “myeloids” (Figure 2E) repeated the UMAP procedure with each subset, which comprised 39 subpopulations in total. There were 13 epithelial cell types (smoker: 24,084 cells, never-smoker: 53,754 cells; Supplementary Figure S4), 7 fibroblastic cell types (smoker: 3,081 cells, never-smoker: 1,592 cells; Supplementary Figure S5), 5 endothelial cell types (smoker: 7,600 cells, never-smoker: 3,783 cells; Supplementary Figure S6), 6 lymphoid cell types (smoker: 25,331 cells, never-smoker: 10,929 cells; Supplementary Figure S7), and 8 myeloid cell types (smoker: 55,398 cells, never-smoker: 40,824 cells; Supplementary Figure S8).

**Figure 2.**
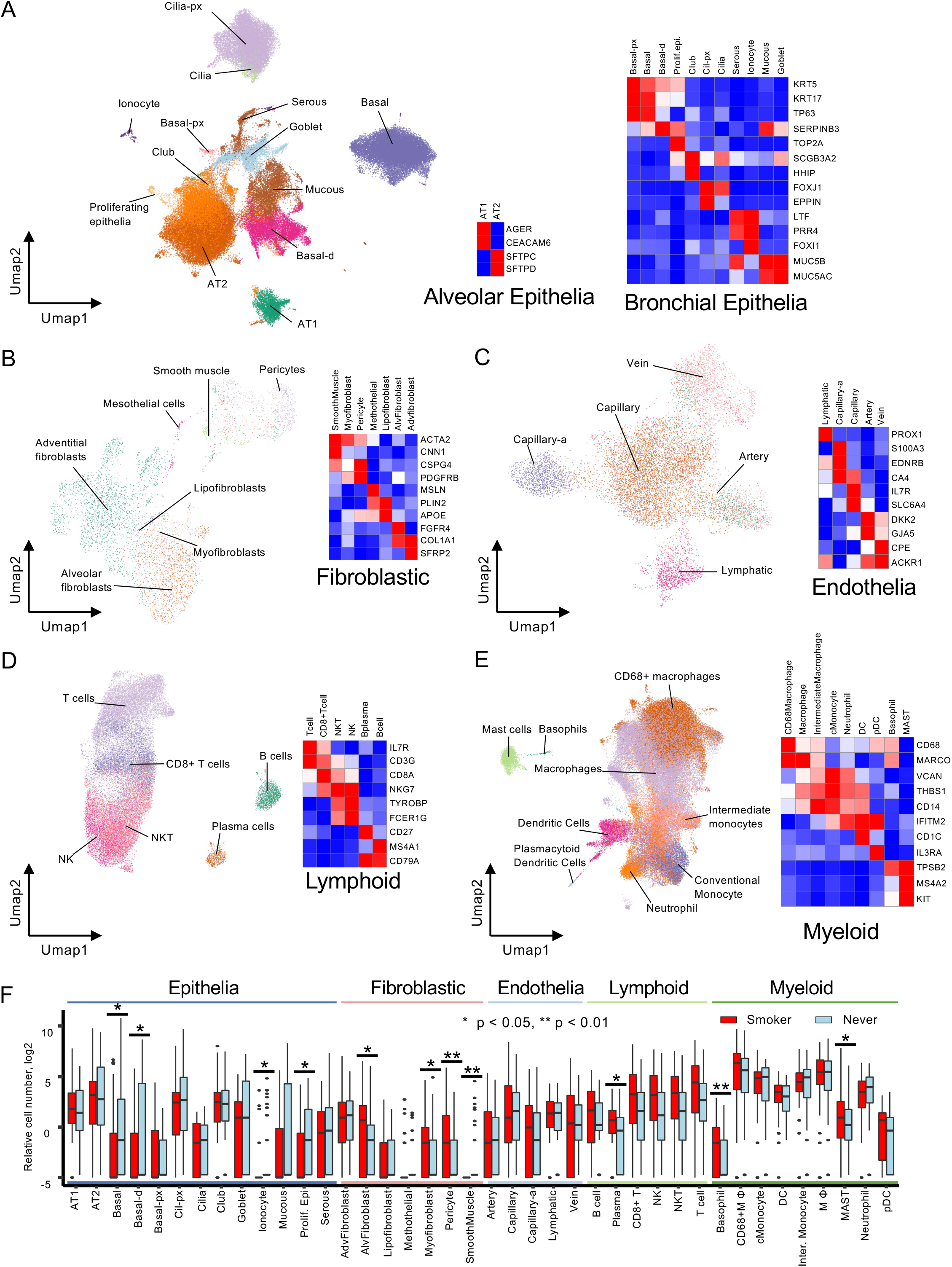
Cell type classification of the integrated lung dataset. UMAP plots for each cell type cluster. The UMAP plot of the cigarette smoking lung atlas was divided into 5 UMAP plots based on the cell type clusters. Heatmaps of selected marker genes in each cell type cluster. Each cluster was defined according to marker expression profiles. **A.** epithelia: 13 clusters, **B.** fibroblasts: 7 clusters. **C.** endothelial cells: 5 clusters. **D.** lymphoid cells: 6 clusters. **E.** myeloid cells: 9 clusters. F. Relative cell number plots between smokers and never-smokers in 39 cell types. Welch’s t test, * p < 0.05. Blue line: epithelial cell types, pink line: fibroblastic cell types, light blue line: endothelial cell types, light green line: lymphoid cell types, and green line: myeloid cell types.

Cigarette smoking is known to induce alterations in cell populations in the lungs. For example, the number of basal linage cells decreased [36], and the number of basophils increased [37] in smoking lungs. The atlas showed differences in the numbers of 39 cell subpopulations by smoking states (Figure 2F). Evidently, the cell numbers of basal, basal-differentiated (d), ionocyte, and proliferating epithelia clusters significantly decreased. Previous bulk studies have reported that the number of bronchial epithelial cells is altered by smoking [9, 36, 38]. Consistent with these reports, our data confirmed that smoking had a devastating effect on epithelial cells in the bronchus and bronchiole. The integrated dataset confirmed the increase in basophil cell number with smoking. We also examined the cell cycle in each cell cluster. The cell cycle indices in each subpopulation were not obviously changed between the smoking and never-smoking groups (Supplementary Figure S9A and B).

### VARIED analysis visualized variations in epithelial populations and basophils by smoking states

Cigarette smoking is the highest risk factor for carcinogenesis of squamous carcinoma in the bronchia and trachea of the lung [2, 5]. To comprehensively understand the effects of smoking in the lung, we developed VARIED (<u>V</u>isualized <u>A</u>lgorithms of <u>R</u>elationships In <u>E</u>xpressional <u>D</u>iversity) analysis to quantify the alteration in gene expressional diversity. VARIED analysis is based on the network centrality of a correlational network with graph theory in each single cell [39, 40]. When the random sampling number of the cells is over 100 cells, the medians of VARIED converged to a certain value (Supplementary Figure S10). Therefore, VARIED analysis needed over 100 cells to calculate the robust centralities. In this study, since all clusters have over 100 cells, we subjected VARIED analysis into the clusters of smoker and never-smoker. The differences in the centrality between smokers and never-smokers represent the alteration of gene expressional diversity in each cell cluster (Figure 3A). VARIED analysis revealed greater diversity in epithelial clusters, suggesting that cigarette smoking primarily perturbed epithelial populations, particularly in the bronchia and trachea (Figure 3B and 3C). These data are consistent with the fact that epithelial cells, located at the bronchia, are considered to be the origin of lung squamous carcinoma [41]. Interestingly, the diversity in basophils was also remarkably altered by cigarette smoking. Basophils are known to be activated as a protective immunity against helminths and ticks by expression of cytokines and Immunoglobins [42]. Basophils in the smokers expressed JUND, FOSB, IGHA1, IGHG1, IGHG3, FCGR3A, and S100A8 (Figure 3D), suggesting their activation and IgG production in smoker lungs.

**Figure 3.**
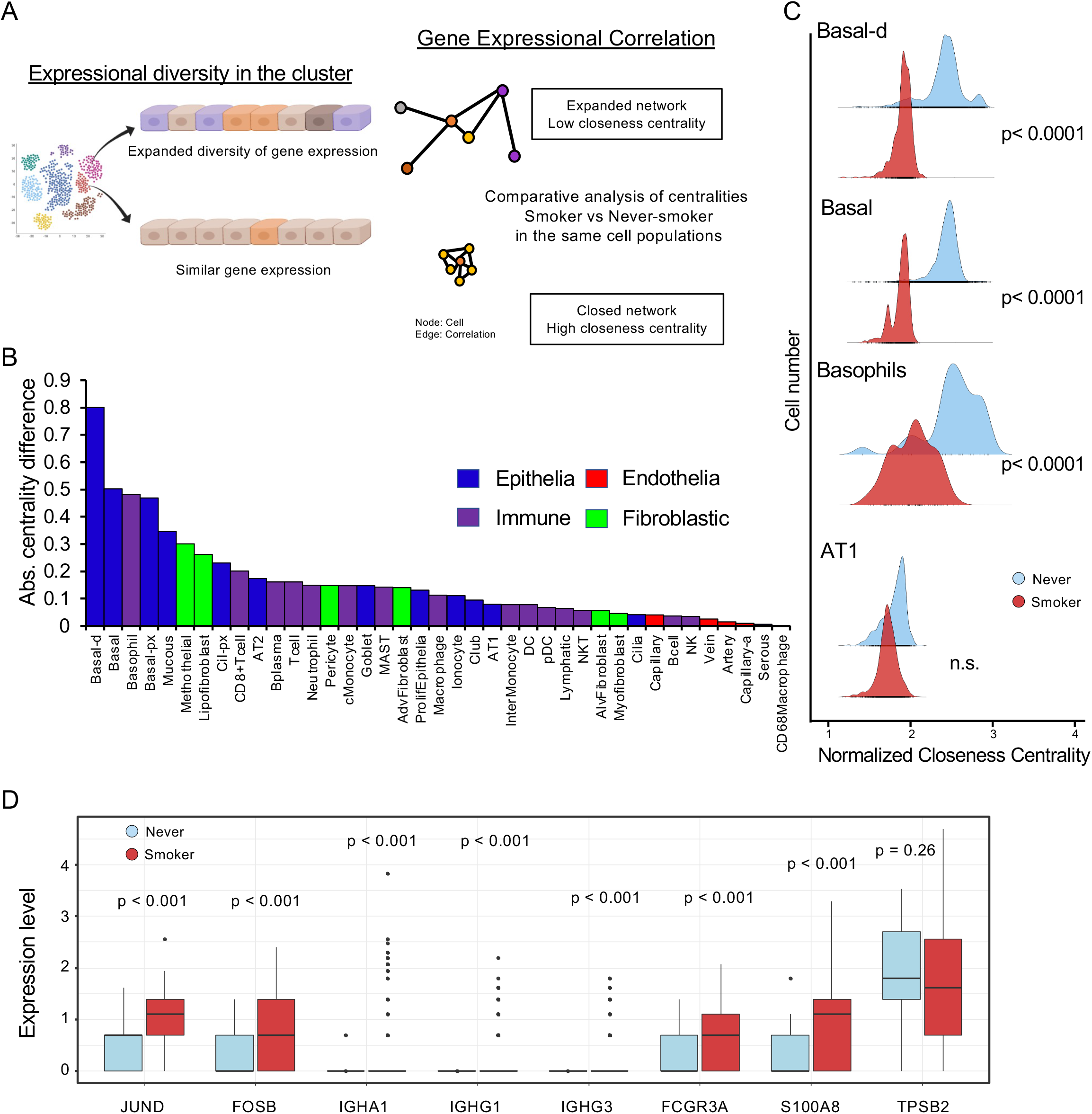
VARIED analysis for cellular variations by smoking states. **A.** Schematic of VARIED (<u>V</u>isualized <u>A</u>lgorithms of <u>R</u>elationships In <u>E</u>xpressional <u>D</u>iversity) analysis for quantifying the alterations in gene expressional diversity between smokers and never-smokers. In each single cell from scRNA-seq, the closeness centrality was calculated in the cell types between smokers and never-smokers. **B.** Plot of absolute values of difference in centrality in each cell type cluster. Blue: epithelia, purple: immune cells, red: endothelia, and green: fibroblasts. **C.** Representative ridge plots for the closeness centrality between smokers and never-smokers. Welch’s t test. **D.** Expression profiles of activation marker gene in the basophil clusters. Box plots of *JUND*, *FOSB*, *IGHA1*, *IGHG1*, *IGHG3*, *FCGR3A*, *S100A8*, and *TPSB2* between smokers and never-smokers in the basophil cluster. Welch’s t test.

To examine the molecular basis for diversity in gene expression, we extracted differentially expressed genes (DEGs) in the basal-d cluster between smokers and never-smokers, focusing on basal-d because this cluster was the most influenced by cigarette smoking (Figure 4, Supplementary Table S3). Enrichment analysis of the DEGs revealed that protein production-related signal, mitochondrial dysfunction pathways were significantly enriched in the smoker basal-px cluster (Figure 4A and B, Supplementary Table S4). Our data indicate that smoking adversely affects bronchial epithelial cells and alters gene expressional diversity in carcinogenesis. The Basal-d cluster in smoker significantly highly expressed ATF3, FOS, and JUN. The VARIED analysis confirmed the early oncogenic events in bronchial and tracheal epithelial cells. In addition, the gene set variation analysis (GSVA) score using the signature genes of smoker basal-d cluster correlated to poor prognosis in lung squamous carcinoma (LUSC) cohorts in the cancer genome atlas (TCGA) (Figure 4D, Supplementary Table S5).

**Figure 4.**
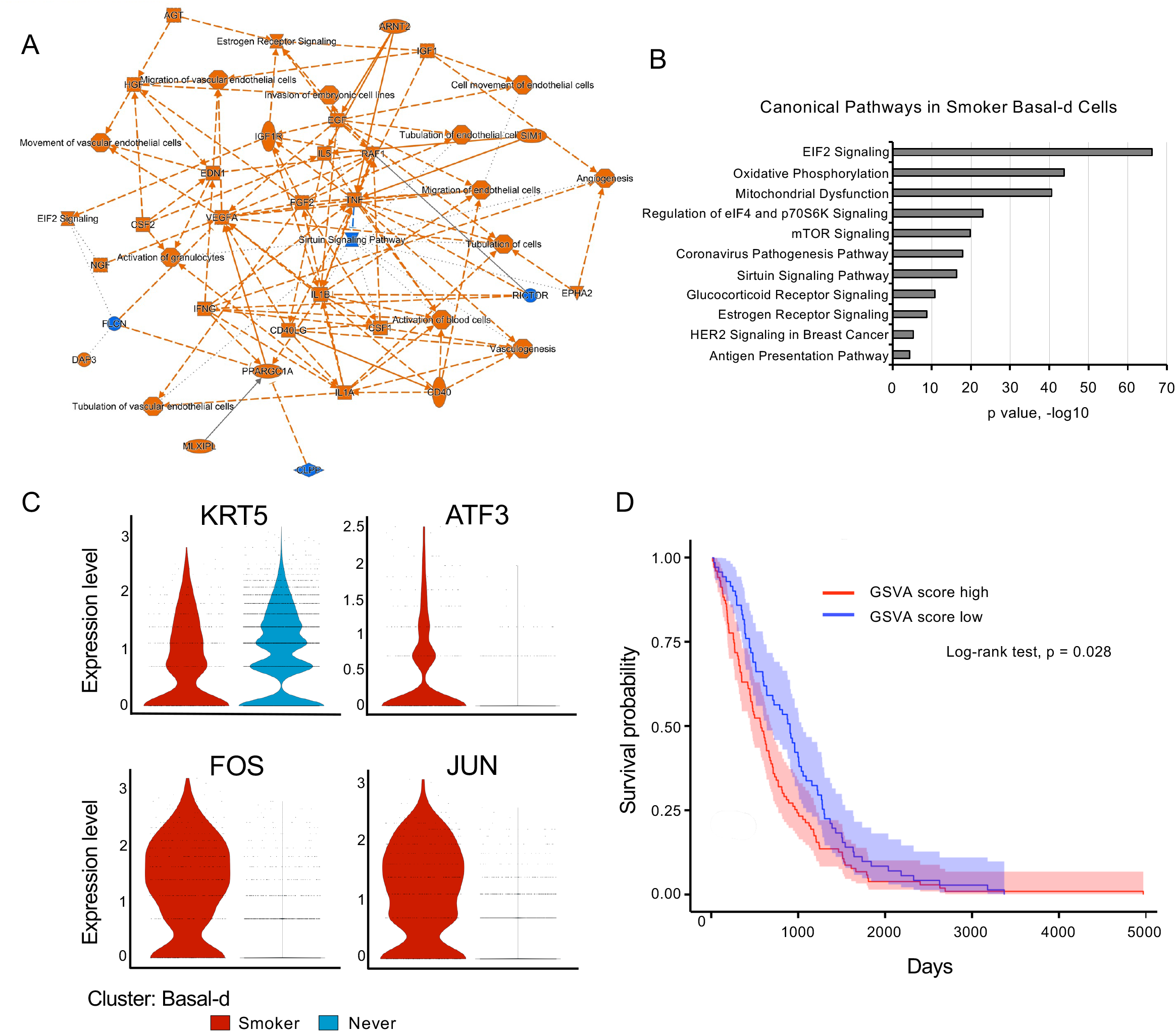
Cancer-related alteration in the Basale-d cluster. Expression profiles of differentially expressional genes (DEGs) in the basal-d clusters between smokers and never-smokers. **A.** Gene and pathway networks of marker genes for the basal-px cluster. The network plot was generated by IPA. **B.** Enrichment analysis of marker genes for the basal-px cluster. Significantly enriched pathways are shown based on IPA data. **C.** Marker expression patterns in the smoker basal-d clusters. Violin plots of *KRT5*, *ATF3*, *FOS*, and *JUN* between smokers and never-smokers. Welch’s t test **D.** Survival analysis of lung squamous carcinoma (LUSC) cohorts in the Cancer Genome Atlas (TCGA) with basal-d smoker signature genes. GSVA scores using smoking basal-d signature genes were calculated by GAVA algorithm. Log-rank test.

### Cigarette smoking affected GWAS-related genes in lung squamous carcinoma

As the cigarette smoking lung atlas provided high-resolution expression data in 39 cell types, we explored gene expression profiles from a genome-wide association study (GWAS) of lung squamous carcinoma with smoking [43]. To identify the expressional patterns and the broad contributions of different lung cell types to squamous carcinoma susceptibility, the expression levels of an average of 92 GWAS genes were examined in all lung cell types (Supplementary Figure S11A). High expression of squamous carcinoma GWAS genes was observed in the specific clusters, and cigarette smoking affected the expression of GWAS-related genes in some clusters. In particular, the expression of *MUC1* was increased in the smoker epithelial clusters (Supplementary Figure S11B), and the expression of *HLA-A* was increased in the smoker myeloid clusters (Supplementary Figure S11C). Mutated *MUC1* has oncogenic roles in carcinogenesis in the human lung [44, 45]. Truncating mutations in *HLA-A* carry a risk of dysregulation of cancer-related pathways [46].

### Cancer-associated alterations induced by smoking

Next, we performed module analysis with cancer-related gene sets, such as senescence, ROS production, IFN signaling, heme metabolism, and epithelial to mesenchymal transition (EMT) genes. The module analysis depicted the alteration of cancer-related events by smoking in each cluster (Figure 5A). Several modules were drastically altered between the smoker and never-smoker groups, such as IFN signaling in endothelial and myeloid clusters; EMT in epithelial, fibroblastic, and endothelial clusters; and mitophagy in lymphoid and myeloid clusters. Because increased expression of EMT module genes in endothelial clusters was observed, we examined the expression of endothelial to mesenchymal transition (EndMT) marker genes (*FN1*, *POSTN*, *VIM*) [17, 47]. These EndMT markers were significantly upregulated, suggesting that smoking induced EndMT in some endothelial clusters (Figure 5B).

**Figure 5.**
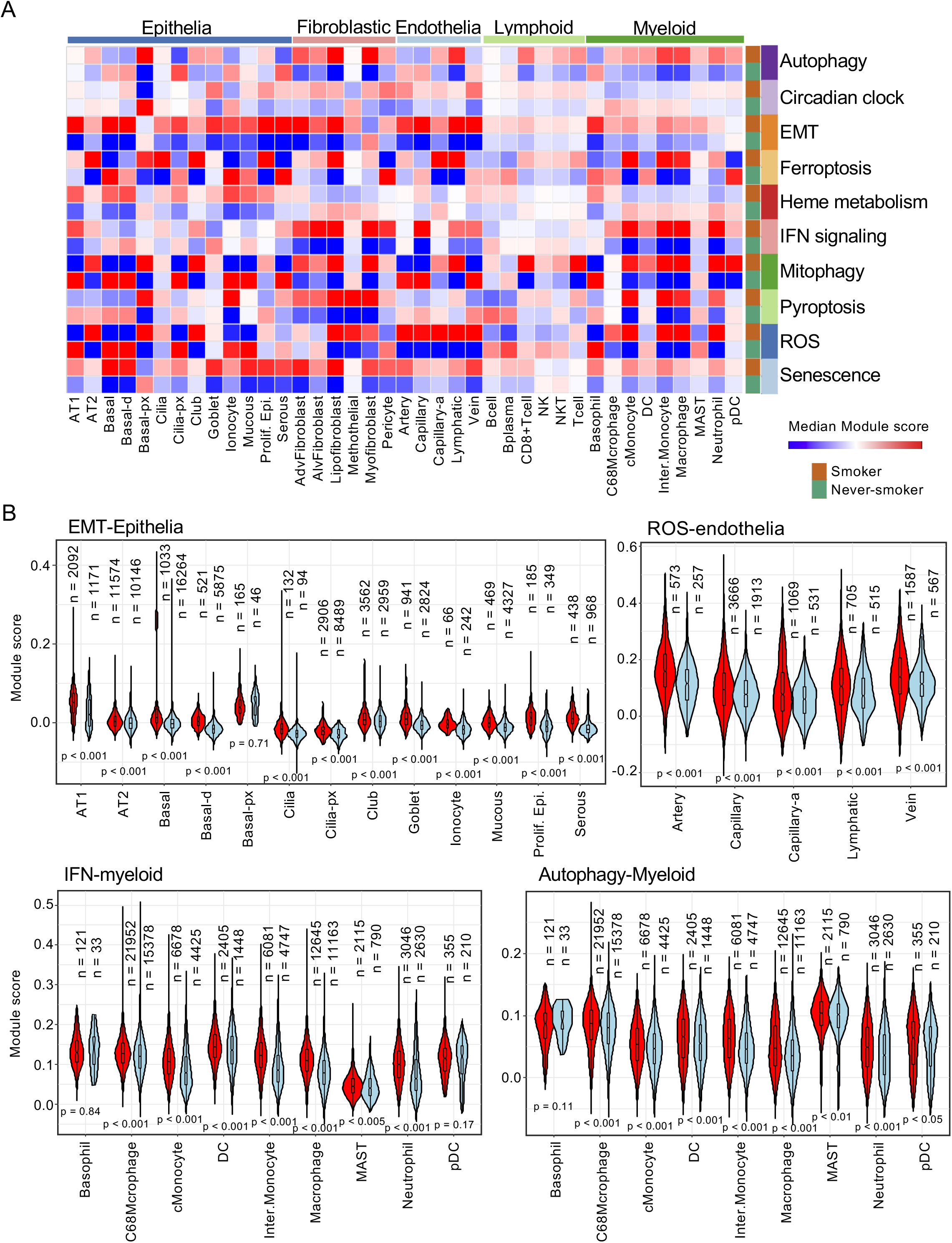
Module analysis of cancer-related pathways in all cell types between smokers and never-smokers. **A.** A heatmap of module analysis between smokers and never-smokers across the cell types. The median module score was calculated for each cell type. The senescence, ROS, pyroptosis, mitophagy, INF signaling, heme metabolism, ferroptosis, EMT circadian clock, and autophagy modules are shown in the heatmap. **B.** Violin plots for analysis of selected modules: EMT, IFN signaling, and heme metabolism. Module analysis for selected cell types is shown. “n” represents the cell number in each cluster. Welch’s t test, p < 0.001.

The module scores of reactive oxygen species (ROS) signaling significantly increased in the endothelial clusters. ROS signaling in endothelia induced inflammatory response and endothelial dysfunction [48, 49]. In addition, Myeloid cells enhanced IFN signaling in smoking lungs (Figure 5B bottom left). Autophagy in immune cells is important for cellular immunity, differentiation, and survival [50]. Autophagy modules increased in immune cells and fibroblastic cells. Finally, increased senescence module scores were broadly observed across most cell types (Supplementary Figure S12), suggesting that smoking induced aging in the lung. The module analysis of the cigarette smoking lung atlas evidently indicated what cell types were influenced by smoking and how smoking affected these cells in the lung.

### Aging-related gene expression in the integrated dataset with cigarette smoking

As the majority of the samples in the atlas had patient age information, we aimed to identify aging-related genes associated with cigarette smoking (Figure 6A). We developed AGED (Aging-related Gene Expression Differences) analysis based on regression analysis with single-cell transcriptome data (see methods). Briefly, by using regression analysis with age and gene expression in the smoker and never-smoker groups, we calculated the differences in slopes (Δ) for all genes in 39 cell clusters (Figure 6B). For selected genes that were obviously changed with advancing age between the smoker and never-smoker groups, the Δ values were plotted as AGED results in a heatmap (Figure 6C). These data showed that the lung surfactant proteins *SFTPC* and *SFTPB* decreased in several epithelial clusters with advancing age in the smoker (Figure 6C and 6D left). These lung surfactant proteins maintain the activation of alveolar macrophages and promote recovery from injuries induced by smoking [51]. Additionally, secretoglobins (*SCGB3A1*, *SCGB3A2*, and *SCGB1A1*) were also decreased in secretory goblet cells and serous cells with advancing age in the smokers. *MALAT1* is a well-known lncRNA in lung cancer, and its expression contributes to malignancy [52, 53]. AGED analysis showed that *MALAT1* expression increased in most cell types with advancing age in smokers (Figure 6C and 6E), suggesting that the oncogenic risk associated with *MALAT1* increased with age. From the module analysis, heme metabolism was dysregulated in the myeloid cells of smoker lung (Figure 5A). High expression of TMSB4X is contributed to poor prognosis and predicts the metastasis in lung carcinoma [53]. TMSB4X increased in most cell types of immune clusters in the smoker lung (Figure 6D right). Finally, the expression level of *FTL* was significantly altered with advancing age in the smokers (Figure 6C). In the cMonocyte clusters, *FTL* significantly decreased with smoking and aging. Collectively, the AGED analysis revealed changes in aging-related gene expression with smoking in each cell cluster.

**Figure 6.**
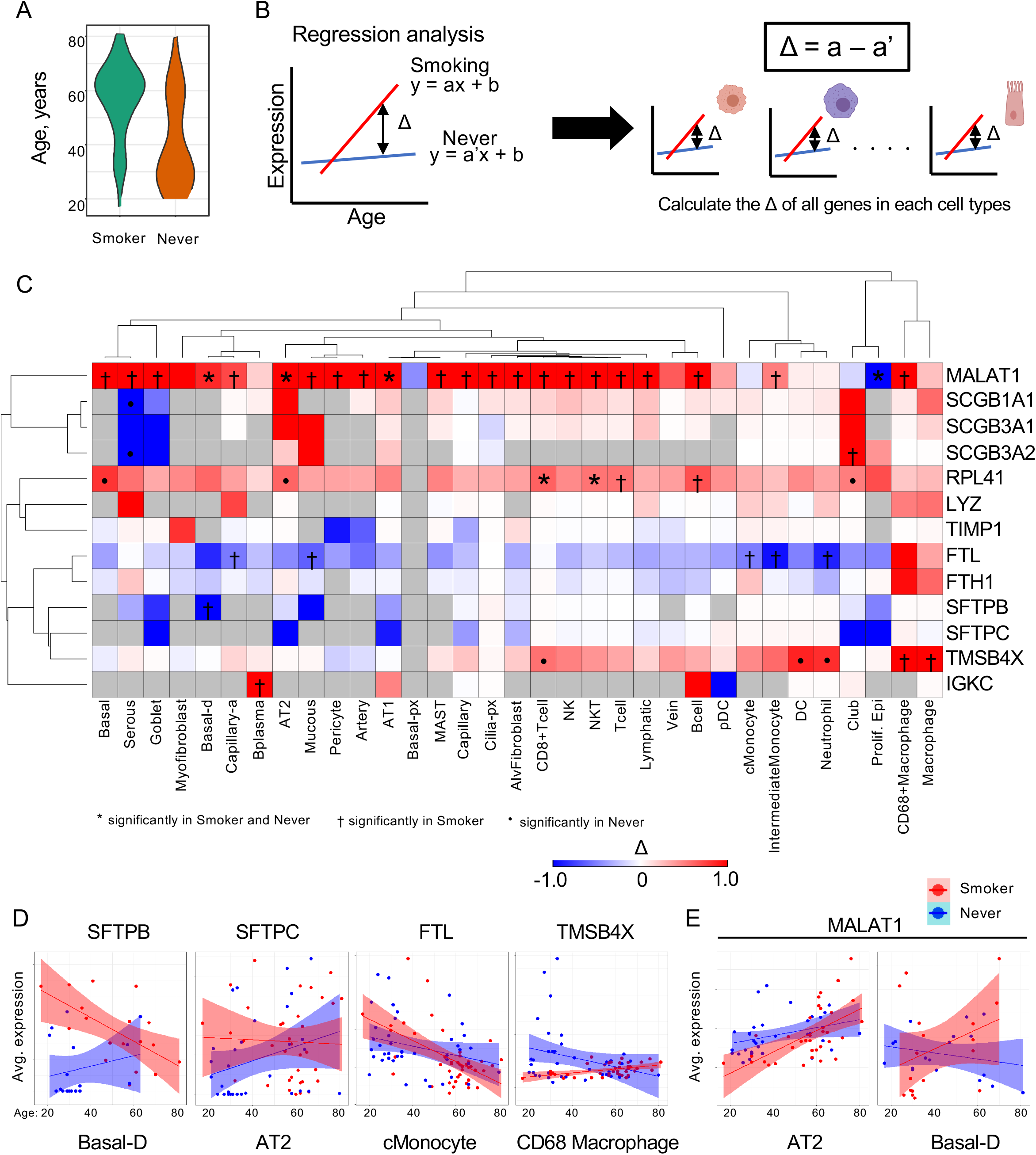
AGED analysis to identify genes related to advancing age and smoking. **A.** Age distributions of the smoker and never-smoker groups. **B.** Schematic of AGED (<u>A</u>ging-related <u>G</u>ene <u>E</u>xpressional <u>D</u>ifferences) analysis for determining gene alterations with advancing age in smokers. Based on regression analysis of single-cell transcriptome data with age, the differences in slopes (Δ) between the smoker and never-smoker groups in 39 cell clusters were calculated for all genes. The Δ values for selected genes were plotted in a heatmap. **C.** Heatmap of AGED analysis results. p value < 0.05: * significant in smokers and never-smokers, † significant in smokers, ‘・’ significant in never-smokers. **D.** Representative correlation plots of *SFTPB* in basal-d cluster, *SFTPC* in AT2 cluster*, FTL* in cMonocyte cluster, and TMSB4X in CD68+Macrophage cluster. **E.** Representative plots of *MALAT1* expression in AT2 and basal-d clusters.

## Discussion

In this study, we presented a human cigarette smoking lung atlas, generated via the meta-analysis of 104 samples from 8 public scRNA-seq datasets. Our integrated smoking atlas confirmed the alteration of gene expression in the lung at single-cell resolution and identified the early oncogenic events induced by cigarette smoking. Additionally, the novel VARIED and AGED analyses revealed cell type and gene expressional diversity with smoking and age.

One of the significant contributions of this study is that the scMeta-analysis of integrated datasets identified expressional diversity in the early phase of lung squamous carcinoma at the single-cell level. In fact, expression analysis following VARIED revealed early oncogenic signaling in epithelial cells, expression changes in GWAS-related genes, and gender-dependent alterations in the smoking lung. In previous studies of the effects of smoking, genetic mutations in oncogenes and tumor suppressor genes were discovered [54-56]. Bronchial epithelial cells from smokers have mutations in *TP53*, *NOTCH1*, *FAT1*, *CHEK2*, *PTEN*, *ARID1A* and other genes [54]. Our atlas showed that survival AKT-mTOR signaling, mitochondrial dysregulation, and sirtuin signaling pathways were altered in bronchial basal cells by smoking (Supplementary Table S4). Mutations in *PTEN* contribute to the activation of AKT-mTOR signaling [57]. *FAT1* controls mitochondrial functions [58], and its mutations induce the dysregulation of mitochondria. Additionally, cigarette smoking promotes lung carcinogenesis by IKKβ- and JNK-dependent inflammation [59]. DEGs analysis of basal-d clusters indicated that *ATF3*, *JUN* and *FOS* expression levels were increased in the smoker basal-d cluster (Supplementary Table S3). High expression of ATF3 expression contributes to tumor malignancy in lung cancer [60]. Our module analysis showed enhancement of inflammatory signaling in the myeloid, fibroblastic, endothelia, and epithelial clusters. The integrated dataset confirmed the signaling related to genetic mutations induced by smoking.

The first scMeta-analysis was performed to investigate severe acute respiratory syndrome coronavirus 2 (SARS-CoV-2)-related genes by The Human Cell Atlas Lung Biological Network [14]. Further scMeta-analyses were reported for endothelial cells in the human and mouse lung [15] and liver-specific immune cells [16], which revealed the alteration of cell populations and expressional heterogeneity with single-cell resolution. Additionally, the study of pan-cancer scRNA-seq cohorts revealed heterogeneity in tumor-infiltrating myeloid cell composition and the functions of cancer-specific myeloid cells [18]. scMeta-analysis is a powerful tool and strategy to overcome the problem of sample bias in small clinical cohorts. Additionally, our integrated dataset enabled us to perform single-cell analysis linked with clinical information in meta-cohorts such as AGED analysis, which identified aging-related gene expression with single-cell resolution. Furthermore, it revealed correlations in the alterations of gene expression associated with smoking and aging. Further scMeta-analyses incorporating additional clinical information will be helpful for understanding homeostasis and diseases.

Our study has limitations. First, differences in the tissue sampling and single-cell isolation methods generated bias in the cell populations used in this study. This bias could not be completely removed by computational normalization. In fact, our integrated datasets showed the differences in cell subpopulations in each dataset (Supplementary Figure S2B). Next, clinical information such as smoking states, gender, and age depended on the collection in the primary studies. The atlas has only a simple classification: smoker or never-smoker; we could not consider detailed smoking information such as the amount of smoking, years of smoking, and Brinkman index (Supplementary Tables S1 and S2). Additionally, patient age was significantly different between the smoker and never-smoker populations (Figure 6A). Moreover, clinical information such as age and gender was not available for all datasets. In the future, it will be necessary to expand the integrated dataset following the publication of new appropriate datasets for a more robust analysis.

The integrated dataset presented herein contributed to the characterization of the alterations caused by cigarette smoking that are related to carcinogenesis of lung squamous carcinoma. However, lung cancer also develops in never-smokers, in whom lung adenocarcinoma is predominant [5, 6]. scMeta-analysis focused on lung adenocarcinoma in different clinical states has the potential to reveal the nature of genetic carcinogenesis. As a future study, the integration of scRNA-seq data from normal lungs (never-smokers) and lung adenocarcinoma could be a feasible approach to discover the mechanism of carcinogenesis and elucidate the cellular diversity in lung adenocarcinoma. In addition, clinical scRNA-seq and scMeta-analysis will be powerful tools in combination with data from pan-cancer multiomics analyses, such as those in TCGA [32, 61, 62]. Therefore, the integration of scMeta-analysis data with clinical and omics data paves the way for an in-depth understanding of the nature of cancer.

## Acknowledgments

We are grateful to all members of the lab for stimulating discussions during the preparation of this manuscript.

## Author contributions

J.N. and Y.Y. conceived and designed the study. J.N. performed the data analysis and construction of the datasets. J.N. and Y.Y. wrote the manuscript. Y.Y. supervised this project. All authors reviewed and edited the manuscript.

## Funding

This work was supported by Project for MEXT KAKENHI (Grant-in-Aid for Scientific Research (B); grant number: 21H02721, Grant-in-Aid for JSPS Fellows: 20J01794, Grant-in-Aid for Early-Carrier Scientists: 21K15562), GSK Japan Research Grant, and Tokyo Biochemical Research Foundation Research Grant.

## Data availability

scMeta-analysis data were available to the NCBI GEO database and EGA database. Detailed information is shown in Supplementary Table 1.

## Competing Interests statement

The authors have declared that no conflict of interest exists.

## Supplemental Figure legends

**Supplementary Figure S1.**
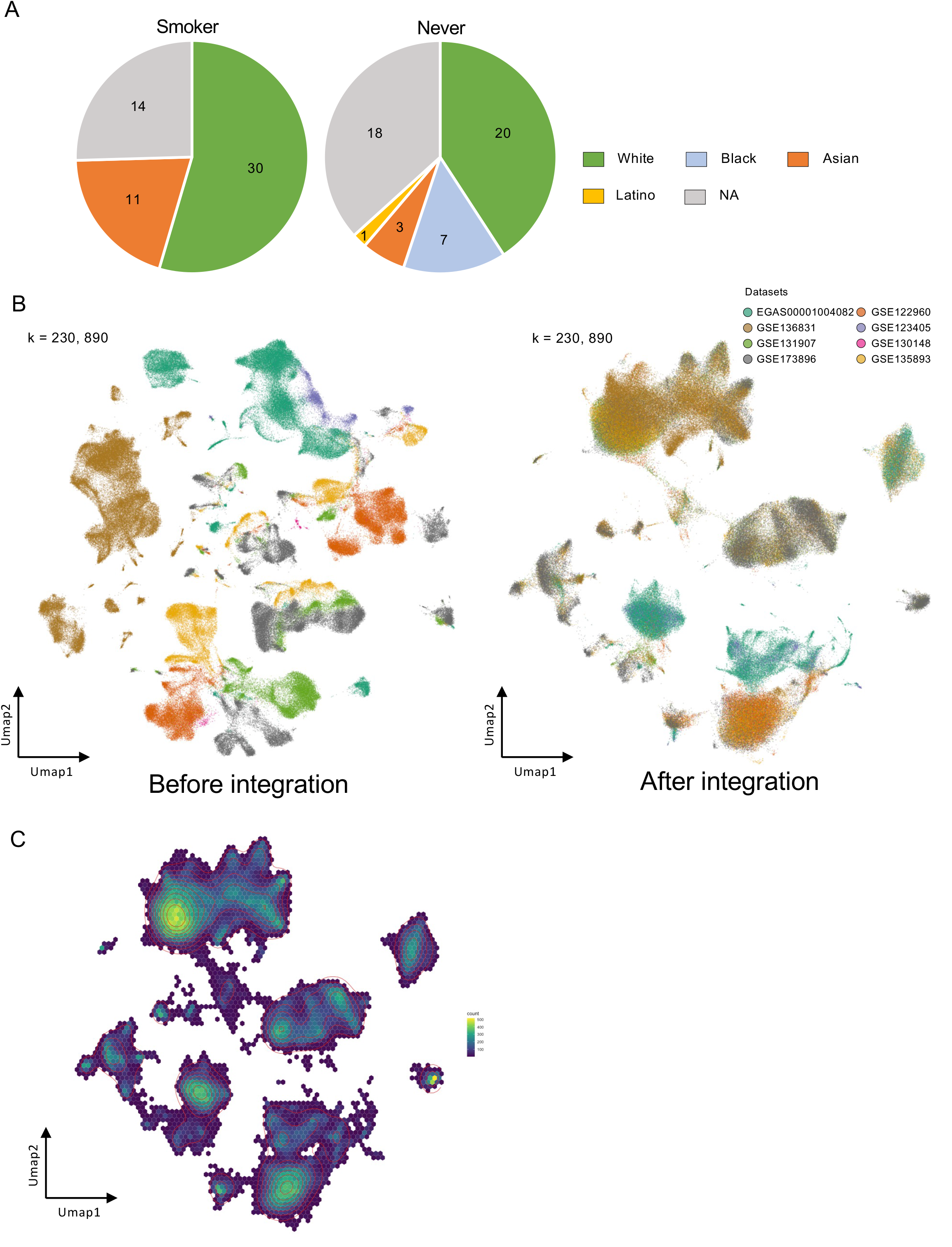
Establishment of the integrated lung scRNA-seq dataset with cigarette smoking states. **A.** The racial distributions of the smoker and never-smoker groups. **B.** A UMAP plots of before harmony integration and after harmony integration of 8 publicly datasets. **C.** A density UMAP plot of the integrated lung dataset.

**Supplementary Figure S2.**
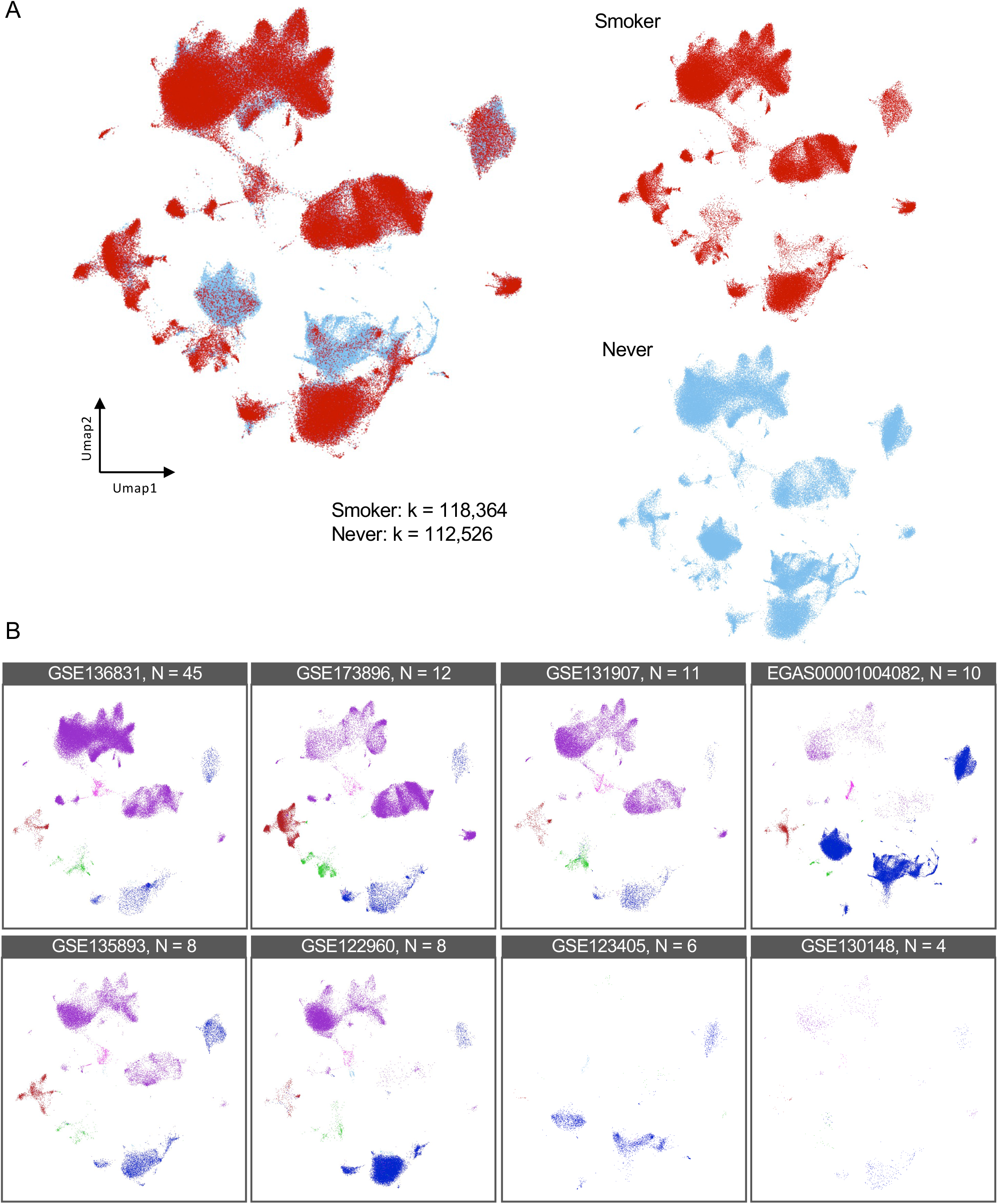
Detailed information of the integrated lung scRNA-seq dataset. **A.** UMAP plot of the integrated lung dataset with smoker/never-smoker information. **B.** Individual UMAP plots for each of 8 publicly available datasets. Blue: epithelia, purple: immune cells, red: endothelia, green: fibroblasts, pink: proliferating immune cells, and light blue: proliferating epithelia.

**Supplementary Figure S3.**
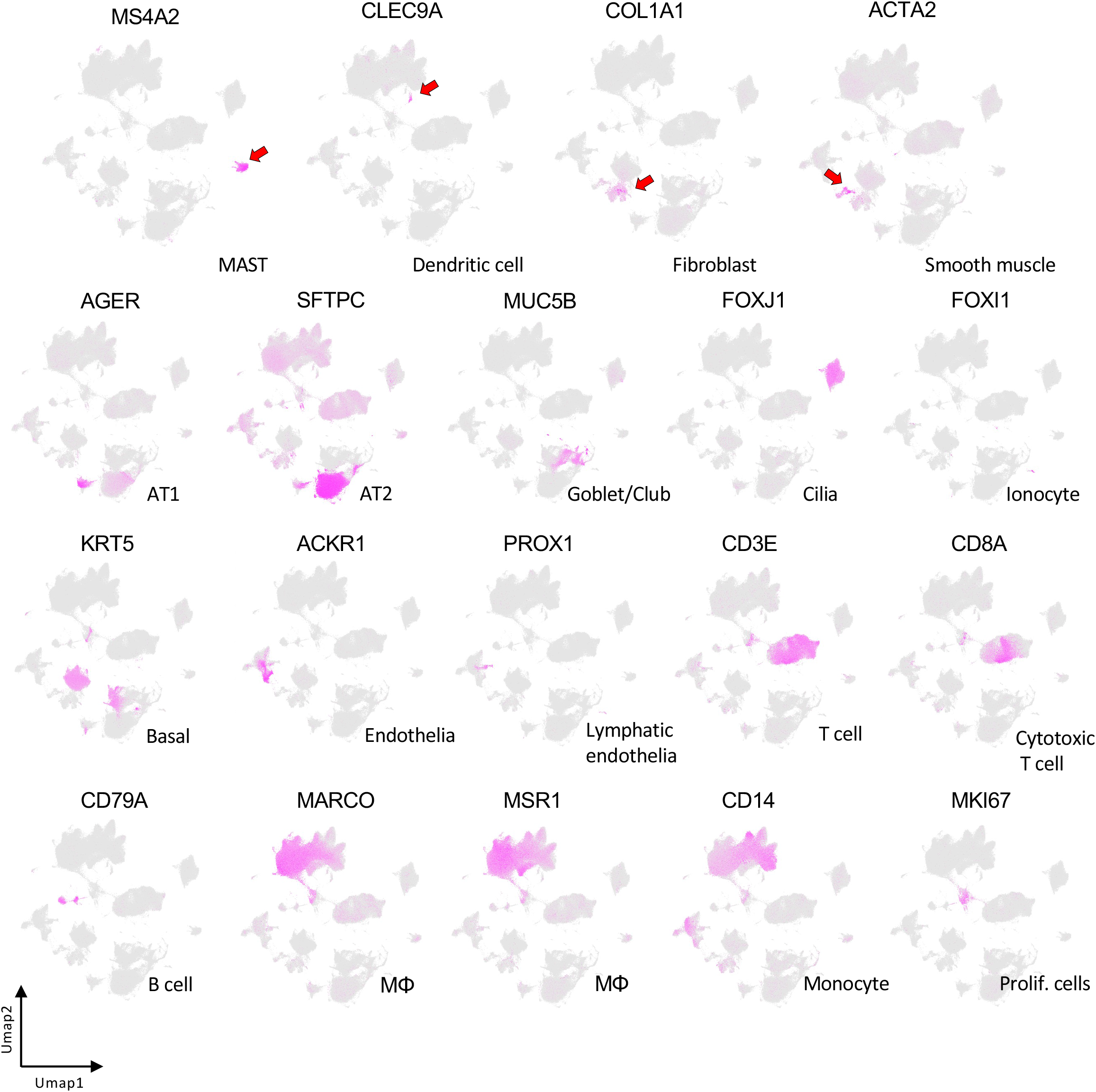
UMAP plots for selected marker genes.

**Supplementary Figure S4.**
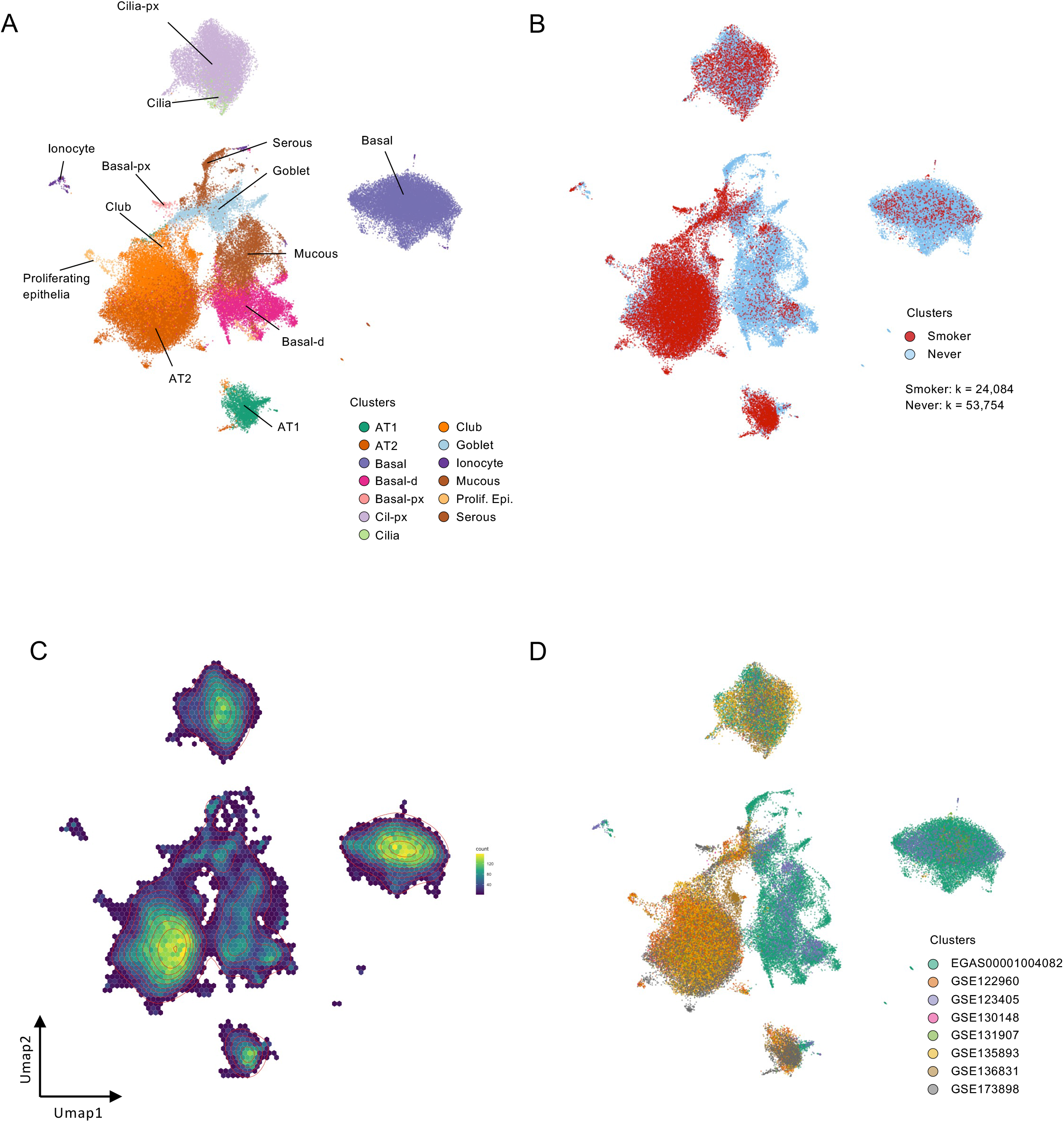
Epithelial cell analysis of smoker and never-smoker lungs. **A.** UMAP plot of 77,838 epithelial cells and proliferating epithelial cells from the UMAP shown in Figure 1D. The dots are labeled by cell type as identified by marker expression profiles. Twelve distinct clusters were identified. **B.** UMAP plot with sample states. Smoker: k = 24,084; never-smoker: k = 53,754. **C.** Density UMAP plot of epithelial cell clusters. **D.** UMAP plot of epithelial cell clusters marked by dataset.

**Supplementary Figure S5.**
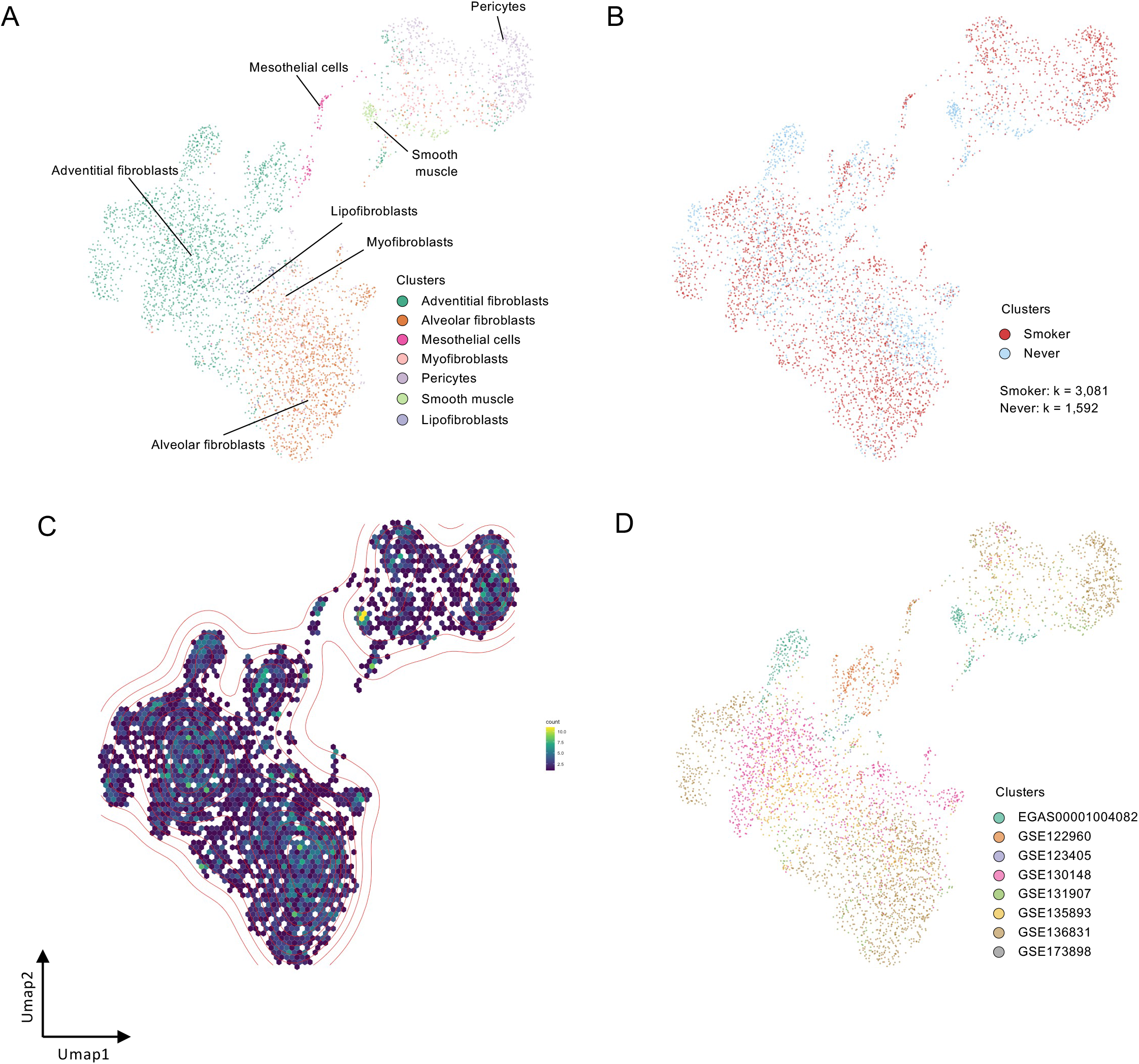
Fibroblast analysis of smoker and never-smoker lungs. **A.** UMAP plot of 4,673 fibroblasts from the UMAP shown in Figure 1B. The dots are labeled by cell type as identified by marker expression profiles. Seven distinct clusters were identified. **B.** UMAP plot with sample states. Smoker: k = 3,081; never-smoker: k = 1,592. **C.** Density UMAP plot of fibroblastic cell clusters. **D.** UMAP plot of fibroblastic cell clusters marked by dataset.

**Supplementary Figure S6.**
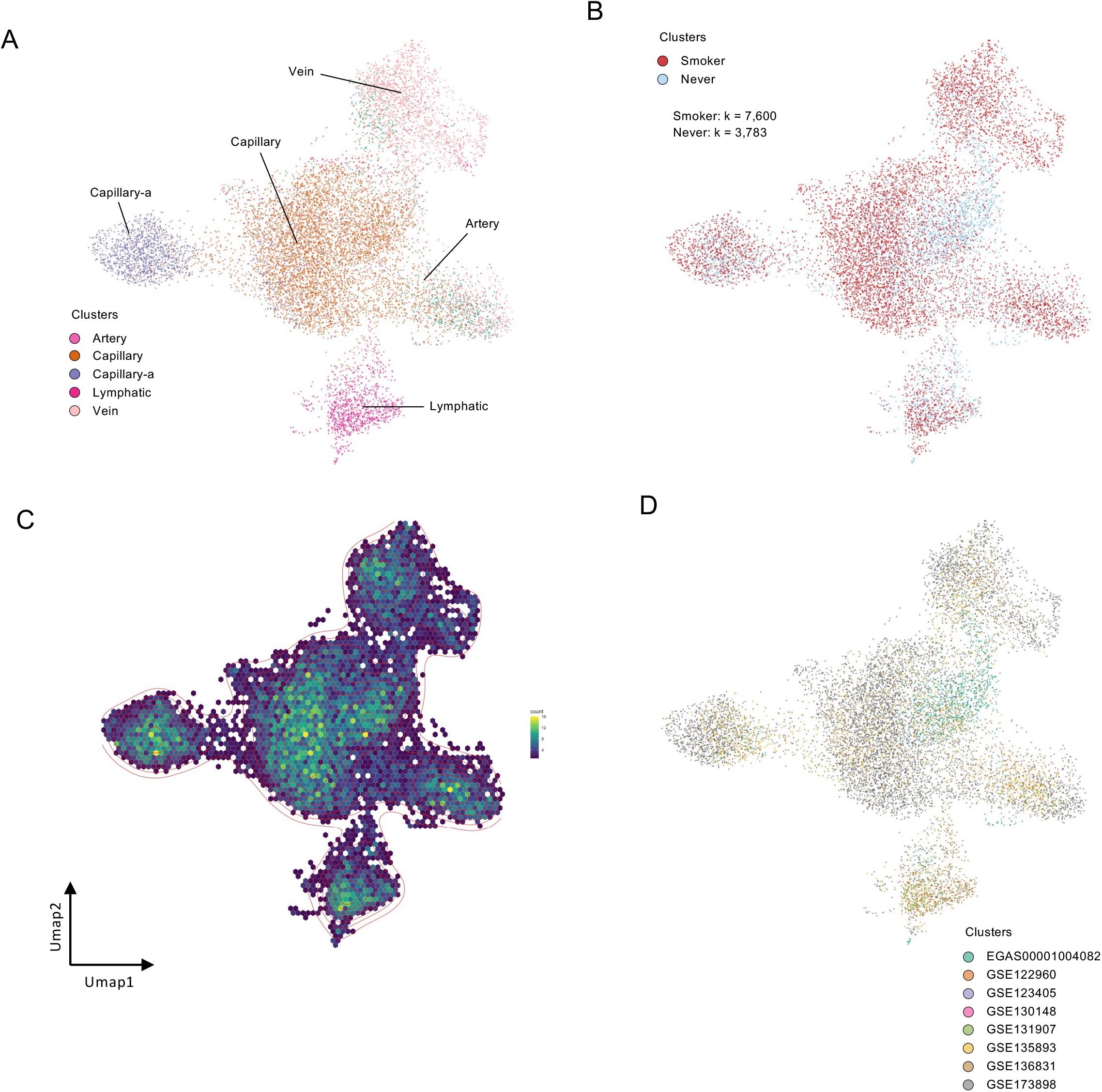
Endothelial cell analysis of smoker and never-smoker lungs. **A.** UMAP plot of 11,383 endothelial cells from the UMAP shown in Figure 1B. The dots are labeled by cell type as identified by marker expression profiles. Seven distinct clusters were identified. **B.** UMAP plot with sample states. Smoker: k = 7,600; never-smoker: k = 3,783. **C.** Density UMAP plot of endothelial cell clusters. **D.** UMAP plot of endothelial cell clusters marked by dataset.

**Supplementary Figure S7.**
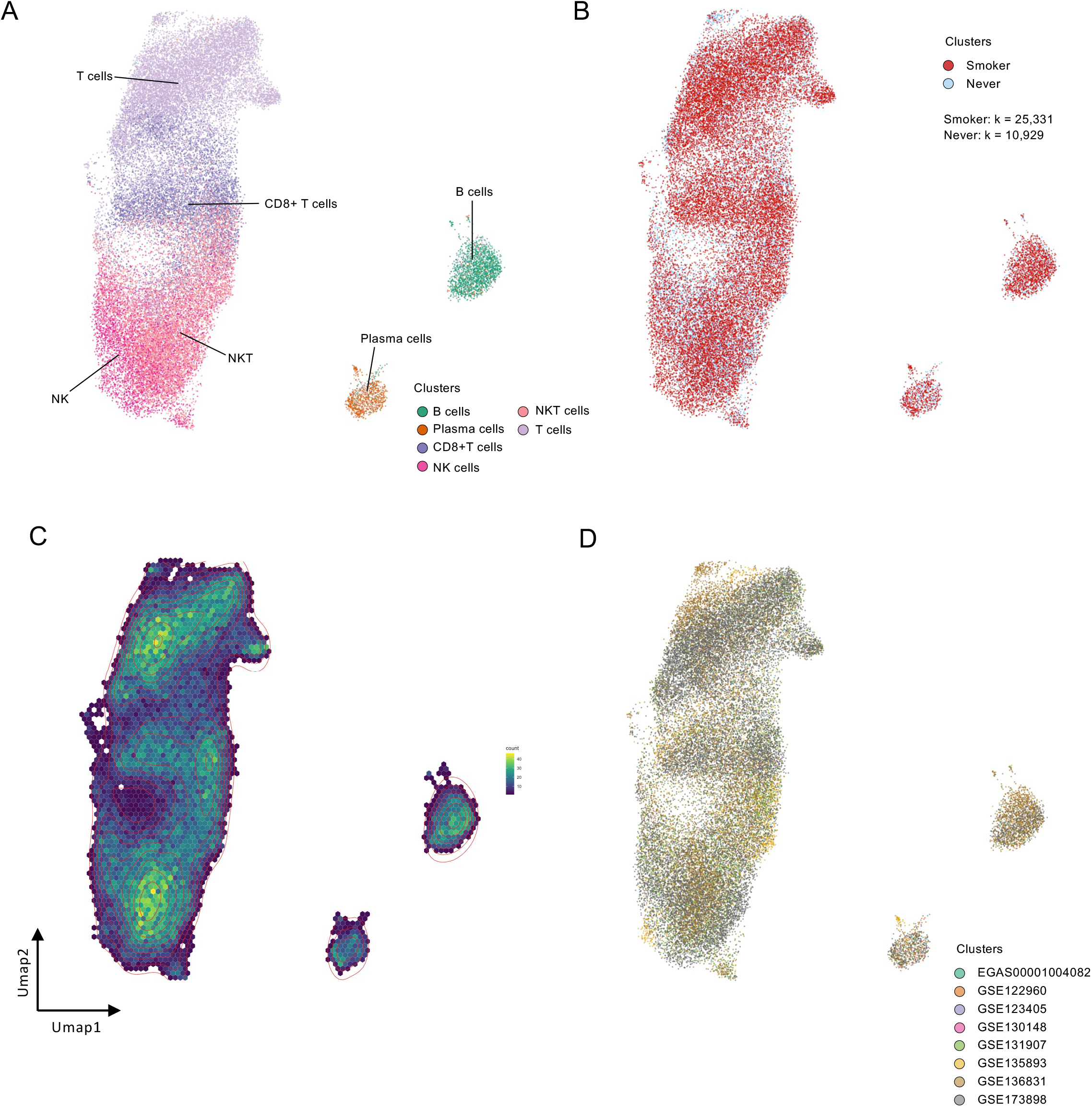
Lymphoid cell analysis of smoker and never-smoker lungs. **A.** UMAP plot of 36,260 lymphoid cells. The dots are labeled by cell type as identified by marker expression profiles. Eight distinct clusters were identified. **B.** UMAP plot with sample states. Smoker: k = 25,331; never-smoker: k = 10,929. **C.** Density UMAP plot of lymphoid cell clusters. **D.** The UMAP plot of lymphoid cell clusters marked by dataset.

**Supplementary Figure S8.**
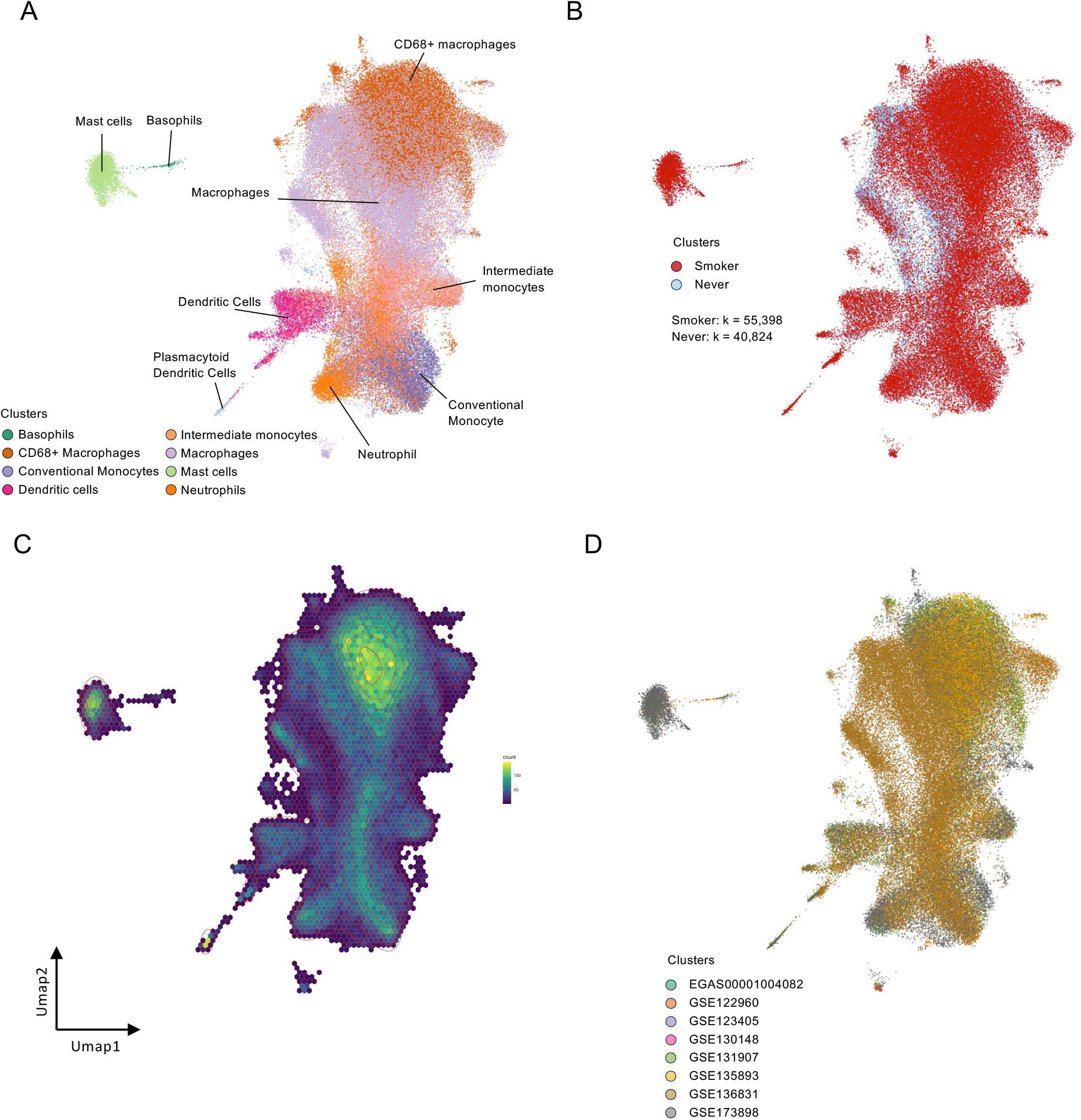
Myeloid cell analysis of smoker and never-smoker lungs. **A.** UMAP plot of 96,222 myeloid cells. The dots are labeled by cell type as identified by marker expression profiles. Eight distinct clusters were identified. **B.** UMAP plot with sample states. Smoker: k = 55,398; never-smoker: k = 40,824. **C.** Density UMAP plot of myeloid cell clusters. **D.** UMAP plot of myeloid cell clusters marked by dataset.

**Supplementary Figure S9.**
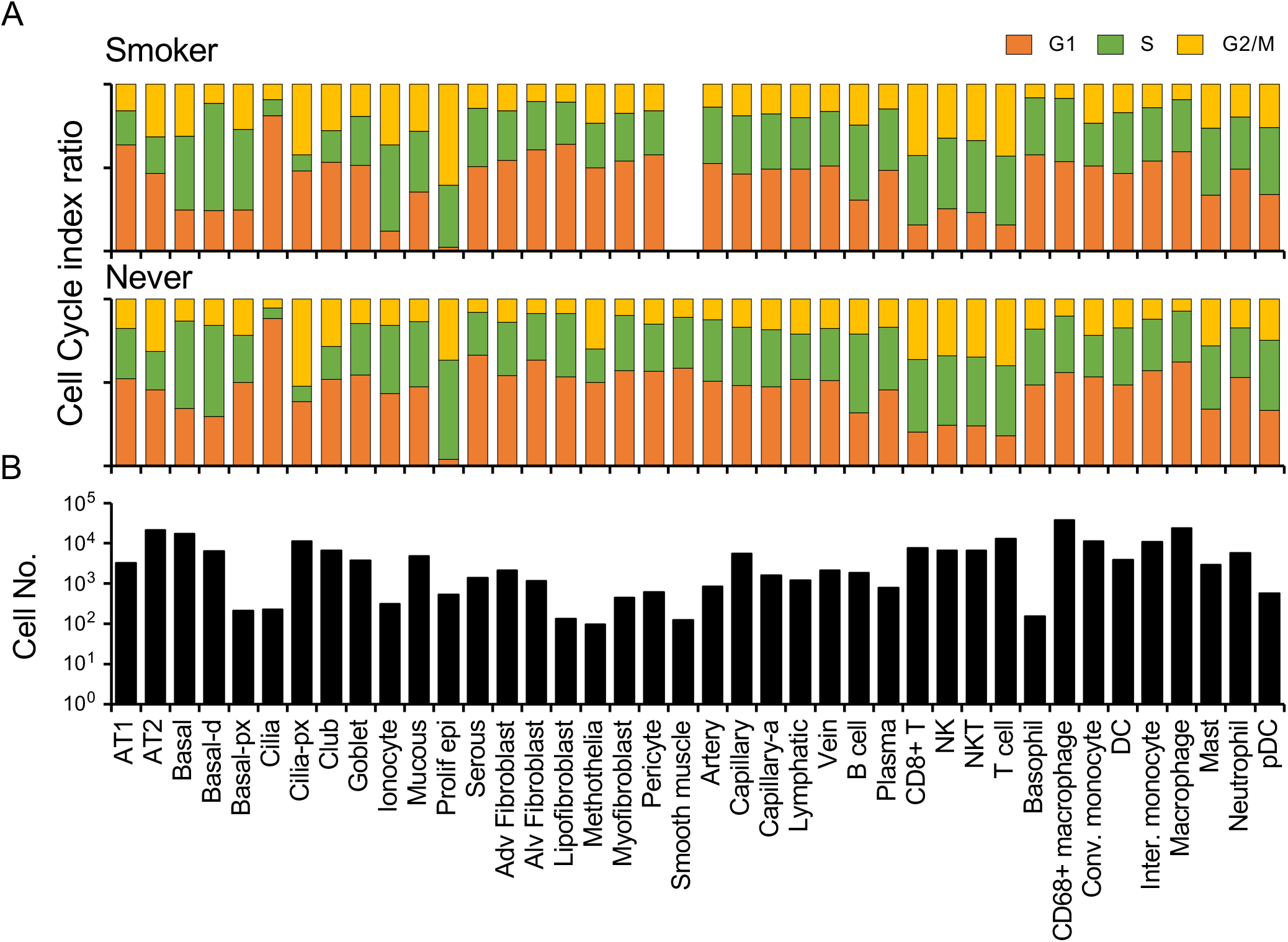
Cell cycle assessment across cell types in the cigarette smoking lung atlas. **A.** Cell cycle phase prediction based on scRNA-seq profiles. G1, S, and G2/M phases are predicted in each cell type. Top: smoker; bottom: never-smoker. **B.** Cell numbers across the cell types.

**Supplementary Figure S10.**
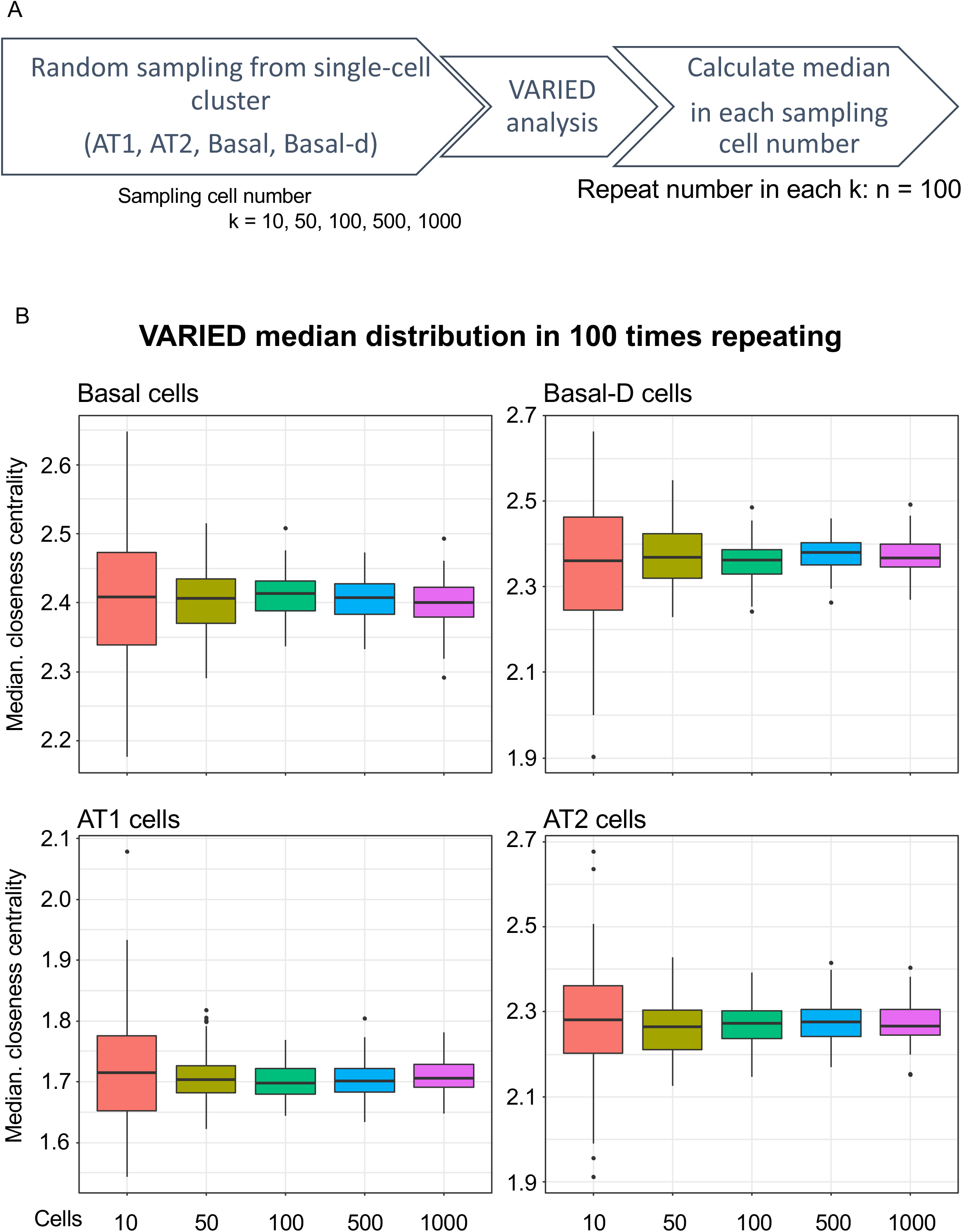
Verifying VARIED analysis by repeating random sampling. **A.** Overview of the accuracy test of VARIED analysis. In first step, the random sampling of cells from the clusters (cell number: k = 10, 50, 100, 500, and 1000) were performed in R. In second step, sampled cells were transformed to correlational network, and performed VARIED analysis. In last step, the median of VARIED score in random sampled cells. It returned to first step. Median calculation was repeated 100 times. **B.** The boxplot showed the distribution of VARIED score median in repeating random sampling test with each sampling cell number.

**Supplementary Figure S11.**
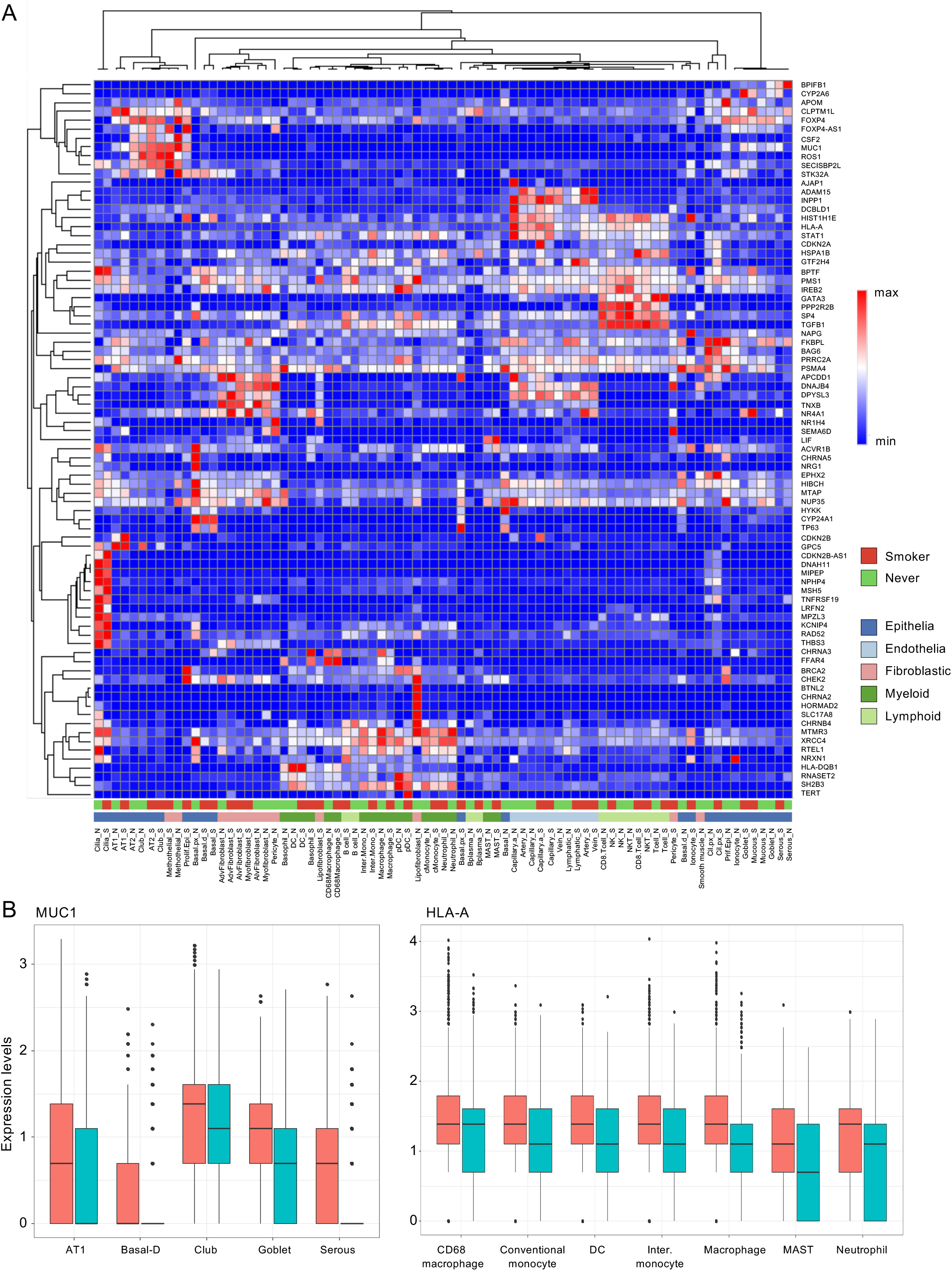
Analysis of GWAS-based squamous cell carcinoma-related genes. **A.** Expression profiles of 92 lung squamous cell carcinoma GWAS genes in all cell types based on the cigarette smoking lung atlas. **B.** MUC1 expression in selected epithelial clusters between the smoker and never-smoker groups. Welch’s t test. **C.** HLA-A expression in selected myeloid clusters between the smoker and never-smoker groups. Welch’s t test.

**Supplementary Figure S12.**
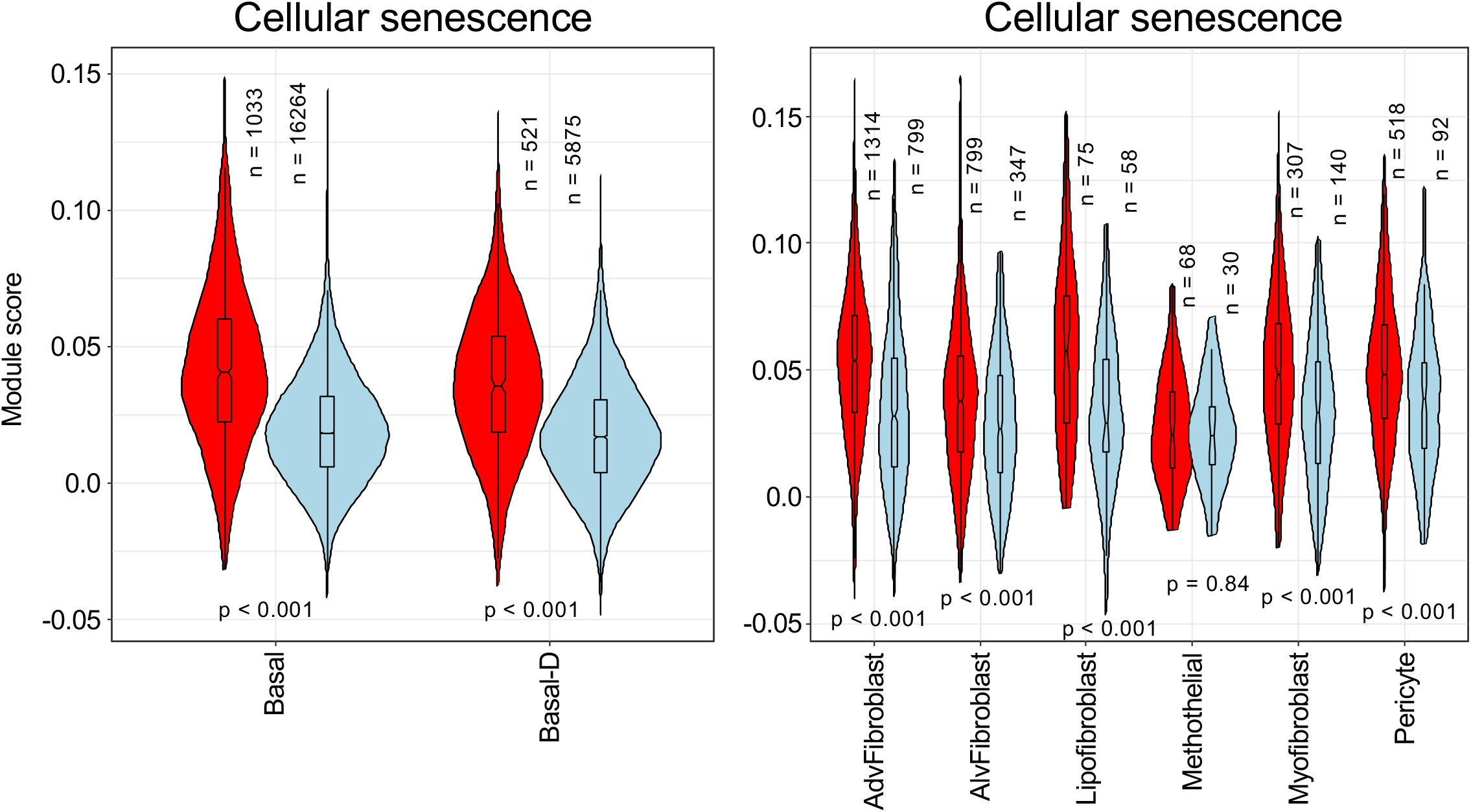
AGED analysis for cellular senescence. A heatmap of AGED analysis results for cellular senescence in basal, basal-d, AdvFibroblast, AlvFibroblast, Lipofibroblast, methothelial, myofibroblast, and pericyte between smokers and never-smokers. “n” represents the cell number in each cluster. Welch’s t test.

## Supplemental Tables

**Supplementary Table S1.** A list of publicly-available 8 datasets for the atlas.

**Supplementary Table S2.** The details of integrated scRNA-seq samples in the atlas.

**Supplementary Table S3.** The DEGs list in basal-d clusters.

**Supplementary Table S4.** IPA canonical pathways in smoker basal-d cluster.

**Supplementary Table S5.** Signature genes of smoking basal-d clusters from DEGs analysis.

## Notes

https://github.com/JunNakayama/scMeta-analysis-of-cigarette-smoking

### Competing Interest Statement

The authors have declared no competing interest.

### Summary of Updates

The atlas was reanalyzed using SCT in Seurat.

## Reference

1. GBD2015Tobacco-Collaborators: Smoking prevalence and attributable disease burden in 195 countries and territories, 1990-2015: a systematic analysis from the Global Burden of Disease Study 2015. Lancet 2017, 389:1885-1906.

2. Stämpfli MR, Anderson GP: How cigarette smoke skews immune responses to promote infection, lung disease and cancer. Nat Rev Immunol 2009, 9:377–384.

3. Lee J, Taneja V, Vassallo R: Cigarette smoking and inflammation: cellular and molecular mechanisms. J Dent Res 2012, 91:142–149.

4. Yanbaeva DG, Dentener MA, Creutzberg EC, Wesseling G, Wouters EF: Systemic effects of smoking. Chest 2007, 131:1557–1566.

5. Sun S, Schiller JH, Gazdar AF: Lung cancer in never smokers--a different disease. Nat Rev Cancer 2007, 7:778–790.

6. Pikor LA, Ramnarine VR, Lam S, Lam WL: Genetic alterations defining NSCLC subtypes and their therapeutic implications. Lung Cancer 2013, 82:179–189.

7. Travaglini KJ, Nabhan AN, Penland L, Sinha R, Gillich A, Sit RV, Chang S, Conley SD, Mori Y, Seita J, et al: A molecular cell atlas of the human lung from single-cell RNA sequencing. Nature 2020, 587:619–625.

8. Goldfarbmuren KC, Jackson ND, Sajuthi SP, Dyjack N, Li KS, Rios CL, Plender EG, Montgomery MT, Everman JL, Bratcher PE, et al: Dissecting the cellular specificity of smoking effects and reconstructing lineages in the human airway epithelium. Nat Commun 2020, 11:2485.

9. Duclos GE, Teixeira VH, Autissier P, Gesthalter YB, Reinders-Luinge MA, Terrano R, Dumas YM, Liu G, Mazzilli SA, Brandsma CA, et al: Characterizing smoking-induced transcriptional heterogeneity in the human bronchial epithelium at single-cell resolution. Sci Adv 2019, 5:eaaw3413.

10. Plasschaert LW, Žilionis R, Choo-Wing R, Savova V, Knehr J, Roma G, Klein AM, Jaffe AB: A single-cell atlas of the airway epithelium reveals the CFTR-rich pulmonary ionocyte. Nature 2018, 560:377–381.

11. Adams TS, Schupp JC, Poli S, Ayaub EA, Neumark N, Ahangari F, Chu SG, Raby BA, DeIuliis G, Januszyk M, et al: Single-cell RNA-seq reveals ectopic and aberrant lung-resident cell populations in idiopathic pulmonary fibrosis. Sci Adv 2020, 6:eaba1983.

12. Habermann AC, Gutierrez AJ, Bui LT, Yahn SL, Winters NI, Calvi CL, Peter L, Chung MI, Taylor CJ, Jetter C, et al: Single-cell RNA sequencing reveals profibrotic roles of distinct epithelial and mesenchymal lineages in pulmonary fibrosis. Sci Adv 2020, 6:eaba1972.

13. Vieira Braga FA, Kar G, Berg M, Carpaij OA, Polanski K, Simon LM, Brouwer S, Gomes T, Hesse L, Jiang J, et al: A cellular census of human lungs identifies novel cell states in health and in asthma. Nat Med 2019, 25:1153–1163.

14. Muus C, Luecken MD, Eraslan G, Sikkema L, Waghray A, Heimberg G, Kobayashi Y, Vaishnav ED, Subramanian A, Smillie C, et al: Single-cell meta-analysis of SARS-CoV-2 entry genes across tissues and demographics. Nat Med 2021, 27:546–559.

15. Schupp JC, Adams TS, Cosme C, Jr., Raredon MSB, Yuan Y, Omote N, Poli S, Chioccioli M, Rose KA, Manning EP, et al: Integrated Single-Cell Atlas of Endothelial Cells of the Human Lung. Circulation 2021, 144:286–302.

16. Rocque B, Barbetta A, Singh P, Goldbeck C, Helou DG, Loh YE, Ung N, Lee J, Akbari O, Emamaullee J: Creation of a Single Cell RNASeq Meta-Atlas to Define Human Liver Immune Homeostasis. Front Immunol 2021, 12:679521.

17. Goveia J, Rohlenova K, Taverna F, Treps L, Conradi LC, Pircher A, Geldhof V, de Rooij L, Kalucka J, Sokol L, et al: An Integrated Gene Expression Landscape Profiling Approach to Identify Lung Tumor Endothelial Cell Heterogeneity and Angiogenic Candidates. Cancer Cell 2020, 37:21–36.e13.

18. Cheng S, Li Z, Gao R, Xing B, Gao Y, Yang Y, Qin S, Zhang L, Ouyang H, Du P, et al: A pan-cancer single-cell transcriptional atlas of tumor infiltrating myeloid cells. Cell 2021, 184:792–809.e723.

19. Xi NM, Li JJ: Benchmarking Computational Doublet-Detection Methods for Single-Cell RNA Sequencing Data. Cell Syst 2021, 12:176–194.e176.

20. Deprez M, Zaragosi LE, Truchi M, Becavin C, Ruiz García S, Arguel MJ, Plaisant M, Magnone V, Lebrigand K, Abelanet S, et al: A Single-Cell Atlas of the Human Healthy Airways. Am J Respir Crit Care Med 2020, 202:1636–1645.

21. Reyfman PA, Walter JM, Joshi N, Anekalla KR, McQuattie-Pimentel AC, Chiu S, Fernandez R, Akbarpour M, Chen CI, Ren Z, et al: Single-Cell Transcriptomic Analysis of Human Lung Provides Insights into the Pathobiology of Pulmonary Fibrosis. Am J Respir Crit Care Med 2019, 199:1517–1536.

22. Zuo WL, Rostami MR, Shenoy SA, LeBlanc MG, Salit J, Strulovici-Barel Y, O’Beirne SL, Kaner RJ, Leopold PL, Mezey JG, et al: Cell-specific expression of lung disease risk-related genes in the human small airway epithelium. Respir Res 2020, 21:200.

23. Watanabe N, Nakayama J, Fujita Y, Mori Y, Kadota T, Shimomura I, Ohtsuka T, Okamoto K, Araya J, Kuwano K, Yamamoto Y: Single-cell Transcriptome Analysis Reveals an Anomalous Epithelial Variation and Ectopic Inflammatory Response in Chronic Obstructive Pulmonary Disease. medRxiv 2020:2020.2012.2003.20242412.

24. Kim N, Kim HK, Lee K, Hong Y, Cho JH, Choi JW, Lee JI, Suh YL, Ku BM, Eum HH, et al: Single-cell RNA sequencing demonstrates the molecular and cellular reprogramming of metastatic lung adenocarcinoma. Nat Commun 2020, 11:2285.

25. Stuart T, Butler A, Hoffman P, Hafemeister C, Papalexi E, Mauck WM, 3rd, Hao Y, Stoeckius M, Smibert P, Satija R: Comprehensive Integration of Single-Cell Data. Cell 2019, 177:1888–1902.e1821.

26. Hafemeister C, Satija R: Normalization and variance stabilization of single-cell RNA-seq data using regularized negative binomial regression. Genome Biol 2019, 20:296.

27. McGinnis CS, Murrow LM, Gartner ZJ: DoubletFinder: Doublet Detection in Single-Cell RNA Sequencing Data Using Artificial Nearest Neighbors. Cell Syst 2019, 8:329–337.e324.

28. Korsunsky I, Millard N, Fan J, Slowikowski K, Zhang F, Wei K, Baglaenko Y, Brenner M, Loh PR, Raychaudhuri S: Fast, sensitive and accurate integration of single-cell data with Harmony. Nat Methods 2019, 16:1289–1296.

29. Tran HTN, Ang KS, Chevrier M, Zhang X, Lee NYS, Goh M, Chen J: A benchmark of batch-effect correction methods for single-cell RNA sequencing data. Genome Biol 2020, 21:12.

30. Nakayama J, Ito E, Fujimoto J, Watanabe S, Semba K: Comparative analysis of gene regulatory networks of highly metastatic breast cancer cells established by orthotopic transplantation and intra-circulation injection. Int J Oncol 2017, 50:497–504.

31. Hänzelmann S, Castelo R, Guinney J: GSVA: gene set variation analysis for microarray and RNA-seq data. BMC Bioinformatics 2013, 14:7.

32. Hoadley KA, Yau C, Hinoue T, Wolf DM, Lazar AJ, Drill E, Shen R, Taylor AM, Cherniack AD, Thorsson V, et al: Cell-of-Origin Patterns Dominate the Molecular Classification of 10,000 Tumors from 33 Types of Cancer. Cell 2018, 173:291–304.e296.

33. Yu G, Wang LG, Han Y, He QY: clusterProfiler: an R package for comparing biological themes among gene clusters. Omics 2012, 16:284–287.

34. Yu G, He QY: ReactomePA: an R/Bioconductor package for reactome pathway analysis and visualization. Mol Biosyst 2016, 12:477–479.

35. Finak G, McDavid A, Yajima M, Deng J, Gersuk V, Shalek AK, Slichter CK, Miller HW, McElrath MJ, Prlic M, et al: MAST: a flexible statistical framework for assessing transcriptional changes and characterizing heterogeneity in single-cell RNA sequencing data. Genome Biol 2015, 16:278.

36. Lumsden AB, McLean A, Lamb D: Goblet and Clara cells of human distal airways: evidence for smoking induced changes in their numbers. Thorax 1984, 39:844-849.

37. Shibata S, Miyake K, Tateishi T, Yoshikawa S, Yamanishi Y, Miyazaki Y, Inase N, Karasuyama H: Basophils trigger emphysema development in a murine model of COPD through IL-4-mediated generation of MMP-12-producing macrophages. Proc Natl Acad Sci U S A 2018, 115:13057–13062.

38. Saetta M, Turato G, Baraldo S, Zanin A, Braccioni F, Mapp CE, Maestrelli P, Cavallesco G, Papi A, Fabbri LM: Goblet cell hyperplasia and epithelial inflammation in peripheral airways of smokers with both symptoms of chronic bronchitis and chronic airflow limitation. Am J Respir Crit Care Med 2000, 161:1016–1021.

39. Watanabe N, Fujita Y, Nakayama J, Mori Y, Kadota T, Hayashi Y, Shimomura I, Ohtsuka T, Okamoto K, Araya J, et al: Anomalous Epithelial Variations and Ectopic Inflammatory Response in Chronic Obstructive Pulmonary Disease. Am J Respir Cell Mol Biol 2022.

40. Tokura M, Nakayama J, Prieto-Vila M, Shiino S, Yoshida M, Yamamoto T, Watanabe N, Takayama S, Suzuki Y, Okamoto K, et al: Single-Cell Transcriptome Profiling Reveals Intratumoral Heterogeneity and Molecular Features of Ductal Carcinoma In Situ. Cancer Res 2022, 82:3236–3248.

41. Hanna JM, Onaitis MW: Cell of origin of lung cancer. J Carcinog 2013, 12:6.

43. Voehringer D: Protective and pathological roles of mast cells and basophils. Nature Reviews Immunology 2013, 13:362-375.

43. Bossé Y, Amos CI: A Decade of GWAS Results in Lung Cancer. Cancer Epidemiol Biomarkers Prev 2018, 27:363–379.

44. Nath S, Mukherjee P: MUC1: a multifaceted oncoprotein with a key role in cancer progression. Trends Mol Med 2014, 20:332–342.

45. Kharbanda A, Rajabi H, Jin C, Tchaicha J, Kikuchi E, Wong KK, Kufe D: Targeting the oncogenic MUC1-C protein inhibits mutant EGFR-mediated signaling and survival in non-small cell lung cancer cells. Clin Cancer Res 2014, 20:5423–5434.

46. Wang Y, Gorlova OY, Gorlov IP, Zhu M, Dai J, Albanes D, Lam S, Tardon A, Chen C, Goodman GE, et al: Association Analysis of Driver Gene-Related Genetic Variants Identified Novel Lung Cancer Susceptibility Loci with 20,871 Lung Cancer Cases and 15,971 Controls. Cancer Epidemiol Biomarkers Prev 2020, 29:1423–1429.

47. Zhao G, Lu H, Liu Y, Zhao Y, Zhu T, Garcia-Barrio MT, Chen YE, Zhang J: Single-Cell Transcriptomics Reveals Endothelial Plasticity During Diabetic Atherogenesis. Front Cell Dev Biol 2021, 9:689469.

48. Csiszar A, Podlutsky A, Wolin MS, Losonczy G, Pacher P, Ungvari Z: Oxidative stress and accelerated vascular aging: implications for cigarette smoking. Front Biosci (Landmark Ed) 2009, 14:3128–3144.

49. Emma R, Caruso M, Campagna D, Pulvirenti R, Li Volti G: The Impact of Tobacco Cigarettes, Vaping Products and Tobacco Heating Products on Oxidative Stress. Antioxidants (Basel) 2022, 11.

50. Ma Y, Galluzzi L, Zitvogel L, Kroemer G: Autophagy and cellular immune responses. Immunity 2013, 39:211–227.

51. Zhao CZ, Fang XC, Wang D, Tang FD, Wang XD: Involvement of type II pneumocytes in the pathogenesis of chronic obstructive pulmonary disease. Respir Med 2010, 104:1391–1395.

52. Gutschner T, Hämmerle M, Eissmann M, Hsu J, Kim Y, Hung G, Revenko A, Arun G, Stentrup M, Gross M, et al: The noncoding RNA MALAT1 is a critical regulator of the metastasis phenotype of lung cancer cells. Cancer Res 2013, 73:1180–1189.

53. Ji P, Diederichs S, Wang W, Böing S, Metzger R, Schneider PM, Tidow N, Brandt B, Buerger H, Bulk E, et al: MALAT-1, a novel noncoding RNA, and thymosin beta4 predict metastasis and survival in early-stage non-small cell lung cancer. Oncogene 2003, 22:8031–8041.

54. Yoshida K, Gowers KHC, Lee-Six H, Chandrasekharan DP, Coorens T, Maughan EF, Beal K, Menzies A, Millar FR, Anderson E, et al: Tobacco smoking and somatic mutations in human bronchial epithelium. Nature 2020, 578:266–272.

55. Vaz M, Hwang SY, Kagiampakis I, Phallen J, Patil A, O’Hagan HM, Murphy L, Zahnow CA, Gabrielson E, Velculescu VE, et al: Chronic Cigarette Smoke-Induced Epigenomic Changes Precede Sensitization of Bronchial Epithelial Cells to Single-Step Transformation by KRAS Mutations. Cancer Cell 2017, 32:360–376.e366.

56. Wang X, Ricciuti B, Nguyen T, Li X, Rabin MS, Awad MM, Lin X, Johnson BE, Christiani DC: Association between Smoking History and Tumor Mutation Burden in Advanced Non-Small Cell Lung Cancer. Cancer Res 2021, 81:2566–2573.

57. Hollander MC, Blumenthal GM, Dennis PA: PTEN loss in the continuum of common cancers, rare syndromes and mouse models. Nat Rev Cancer 2011, 11:289–301.

58. Cao LL, Riascos-Bernal DF, Chinnasamy P, Dunaway CM, Hou R, Pujato MA, O’Rourke BP, Miskolci V, Guo L, Hodgson L, et al: Control of mitochondrial function and cell growth by the atypical cadherin Fat1. Nature 2016, 539:575–578.

59. Takahashi H, Ogata H, Nishigaki R, Broide DH, Karin M: Tobacco smoke promotes lung tumorigenesis by triggering IKKbeta- and JNK1-dependent inflammation. Cancer Cell 2010, 17:89–97.

60. Li X, Zhou X, Li Y, Zu L, Pan H, Liu B, Shen W, Fan Y, Zhou Q: Activating transcription factor 3 promotes malignance of lung cancer cells in vitro. Thorac Cancer 2017, 8:181–191.

61. Hutter C, Zenklusen JC: The Cancer Genome Atlas: Creating Lasting Value beyond Its Data. Cell 2018, 173:283–285.

62. Weinstein JN, Collisson EA, Mills GB, Shaw KR, Ozenberger BA, Ellrott K, Shmulevich I, Sander C, Stuart JM: The Cancer Genome Atlas Pan-Cancer analysis project. Nat Genet 2013, 45:1113–1120.

